# Shaping Brain Structure: Genetic and Phylogenetic Axes of Macro Scale Organization of Cortical Thickness

**DOI:** 10.1101/2020.02.10.939561

**Authors:** Sofie L. Valk, Ting Xu, Daniel S. Margulies, Shahrzad Kahrabian Masouleh, Casey Paquola, Alexandros Goulas, Peter Kochunov, Jonathan Smallwood, B.T. Thomas Yeo, Boris C. Bernhardt, Simon B. Eickhoff

**Affiliations:** Institute of Neuroscience and Medicine (INM-7: Brain and Behaviour), Research Centre Jülich, 52425 Jülich, Germany; Institute of Systems Neuroscience, Heinrich Heine University Düsseldorf, 40225 Düsseldorf, Germany; From the Center for the Developing Brain, Child-Mind Institute, New York, USA; Centre National de la Recherche Scientifique, UMR 7225, Frontlab, Institut du Cerveau et de la Moelle Épinière, Paris, France; McConnell Brain Imaging Centre, Montreal Neurological Institute and Hospital, McGill University, Montreal, Canada; Institute of Computational Neuroscience, University Medical Center Hamburg-Eppendorf, Hamburg University, Hamburg, Germany; Maryland Psychiatric Research Center, University of Maryland School of Medicine, Baltimore, Maryland, US; York Neuroimaging Center, University of York, York, United Kingdom; Department of Electrical and Computer Engineering, Centre for Sleep and Cognition, Clinical Imaging Research Centre and N.1 Institute for Health and Memory Networks Program, National University of Singapore, Singapore, Singapore; Athinoula A. Martinos Center for Biomedical Imaging, Massachusetts General Hospital, Charlestown, MA, USA; NUS Graduate School for Integrative Sciences and Engineering, National University of Singapore, Singapore, Singapore

## Abstract

Structural and functional characteristics of the cortex systematically vary along global axes as a function of cytoarchitecture, gene expression, and connectivity. The topology of the cerebral cortex has been proposed to be a prerequisite for the emergence of human cognition and explain both the impact and progression of pathology. However, the neurogenetic origin of these organizational axes in humans remains incompletely understood. To address this gap in the literature our current study assessed macro scale cortical organization through an unsupervised machine learning analysis of cortical thickness covariance patterns and used converging methods to evaluate its genetic basis. In a large-scale sample of twins (n=899) we found structural covariance of thickness to be organized along both an anterior-to-posterior and inferior-to-superior axis. We found that both axes showed a high degree of correspondence in pairs of identical twins, suggesting a strong heritable component in humans. Furthermore, comparing these dimensions in macaques and humans highlighted similar organizational principles in both species demonstrating that these axes of cortical organization are phylogenetically conserved within primate species. Finally, we found that in both humans and macaques the inferior-superior dimension of cortical organization was aligned with the predictions of the dual-origin theory, highlighting the possibility that the macroscale organization of primate brain structure is subject to multiple distinct neurodevelopmental trajectories. Together, our study establishes the genetic basis of natural axes in the cerebral cortex along which structure is organized and so provides important insights into the organization of human cognition that will inform both our understanding of how structure guides function and for the progression of pathology in diseases.

## Introduction

A fundamental question in neuroscience is how the structure of the cortex constrains its function. Over the course of almost a century, numerous studies have shown that the cerebral cortex is organized along dimensions that reflect systematic variations in features of brain structure and function such as laminar differentiation, gene expression, structural and functional connectivity^1–15^. These dimensions have been suggested to reflect the timing of neurogenesis and may relate to the neurogenetic origin of cortical organization ^3, 16^. A potential mechanism for the source of neurogenetic differentiation of brain regions is described by the dual origin theory ^3, 17–21^. This theory conceptualizes cortical areas as emerging from waves of laminar differentiation that spring from the piriform cortex (paleo-cortex) and the hippocampus (archi-cortex). The dual structure might be rooted in heterochronous ontological axes in the developing cortex^16^.

The systematic topological organization of the cerebral cortex has been proposed to reflect an architecture which optimize the balance of externally and internally oriented functioning, which is critical for flexibility of human cognition ^22^. For example, association cortex is located at maximal distance from regions of primary cortex that are functionally specialized for perceiving and acting in the here and now. This increased spatial distance from primary cortex may allow association cortex to take on functions that are only loosely constrained by the immediate environment, allowing internal representations to contribute to cognition and so enhancing the flexibility, and evolutionary fitness of behavior ^22–26^. Accordingly, understanding how the structure of the cortex scaffolds function in a flexible manner requires understanding how macroscale structural features of the organization of the human cortex emerge. Moreover, previous work has implicated macroscale organizational axes of structure and function in the impact and progression of pathology. For example, Parkinson’s and Alzheimer’s disease have been proposed to follow a trajectory, in which underlying anatomical axes determine the sequence in which specific regions and networks are progressively impacted at different disease stages ^27, 28^. Recently, we have been able to show that functional abnormalities in autism spectrum disorder relate to systematic disruptions in large-scale organization of brain function, providing a parsimonious reference frame in which the heterogeneous symptoms of autism spectrum disorder can be understood ^29^.

Although the importance of macroscale axes of cortical organization in cognition and pathology are now recognized, the degree to which these topological features of the cerebral cortex are genetically determined remains incompletely understood. Measured across a population, local brain structure shows marked patterns of covariation across the cerebral cortex, termed ‘structural covariance’. These macro scale patterns in cortical thickness have been linked to both structural and functional connectivity ^30, 31^ and twin studies have shown that thickness covariance between regions is largely due to additive genetic effects ^32, 33^. Recent work shows that inter-regional genetic correlation is determined by two organizational principles: (1) regions are strongly genetically correlated with their counterparts in the opposite cerebral hemisphere ^34, 35^ and (2) regions are highly genetically correlated with geometrically nearby regions ^35^. The local processes that govern the observed distribution of cortical thickness are reasonably well understood. For example, associations between structural and functional connectivity may arise due to shared trophic changes at the synaptic and cellular levels ^36, 37^ and/or reflect coupled expression of genes enriched in supra-granular layers ^38^ that are associated with transcriptomic similarity of local brain regions^39^. Importantly both of these effects converge with postmortem inter-regional correlations of gene expression ^40^. Developmentally, macro scale patterns of cortical thickness mature with age, possibly because of synchronized neurodevelopment ^36, 37^ and the expression of common genetic cues during early cortical development ^41^.

Taken together contemporary theory suggests that (a) macro scale patterns of cortical structure make an important contribution to human cognition and (b) that this is supported by common genetic influences in local areas of cortex. However, we currently lack a clear understanding of how genetic influences contribute to the fundamental organizational principles that underpin the macro scale patterns of cortical thickness seen in humans. Our current study sought to directly examine how genetic influences contribute to the spatial organization of macro scale features of the cortex. We used advanced machine learning methods to construct large-scale organizational gradients that underpin the structural covariance across the cortex. In contrast to clustering-based decompositions of the brain into discrete communities ^42^, cortex-wide gradient mapping techniques describe neural structure and function in a low dimensional space, or, coordinate system, that reflects the macro scale patterns that underpin the observed neural data. We used this approach to describe the structural covariance in humans as well as in non-human primates, and to evaluate whether theses dimensions of variation are genetically determined. In particular, we used a twin-design based on the Human Connectome Young Adult sample (S900) using Sequential Oligogenic Linkage Analysis Routines (www.solar-eclipse-genetics.org; Solar Eclipse 8.4.0.) to evaluate genetic correlation of local cortical thickness across the cortical mantle. In a second analysis we evaluated the phylogenetic basis of macros scale patterns of structural covariance by comparing the large-scale gradients in macaque monkeys (PRIME-DE) ^43^ with those seen in humans. Last, we compared the axes of macro scale organization of cortical thickness in humans and macaques with organizational axes expected based on the theory of dual origin ^3, 17–20^.

Foreshadowing our results, both analyses found evidence that the two main organizational patterns that describe macro scale patterns of cortical thickness were driven by genetic factors. Using a pedigree model to evaluate the genetic correlation of thickness in humans, we found that macro scale patterns of cortical thickness covariance were highly influenced by genetics, especially in prefrontal cortex, highlighting the role of genetics in shaping brain structure in regions functionally associated with complex features of human cognition. We also observed a similar macro scale organization of cortical thickness in humans and macaques, suggesting that these axes are phylogenetically conserved in primates. Moreover, we found an inverse relationship between archi-cortex (hippocampus) and paleo-cortex (olfactory cortex) distance and the inferior-to-superior organization gradient in humans and macaques, aligning covariance topology with the dual origin theory. Together these analyses highlight the important role that genetic processes play in determining the large-scale organization of cortical structure, and so provide an important window into the innate architecture supporting human cognition and a potential model for impact and progression of pathology.

## Results

### Posterior-anterior and inferior-superior axes underlie macro scale organization of cortical thickness

We started our analysis by evaluating the topological organization of structural covariance (Figure 1). We used the mean thickness within 400 parcels ^44^ to create group-level covariance maps based on individual thickness values of participants from the Human Connectome Project (HCP, S900). When computing the macro scale organization of cortical thickness, we controlled for the effects of age, sex, and global thickness. First, we evaluated the average structural covariance as a function of brain network organization ^42^. Strength of structural covariance was stronger between regions within the same functional community than between networks (Figure 1B; Supplementary Table 1).

**Fig 1.**
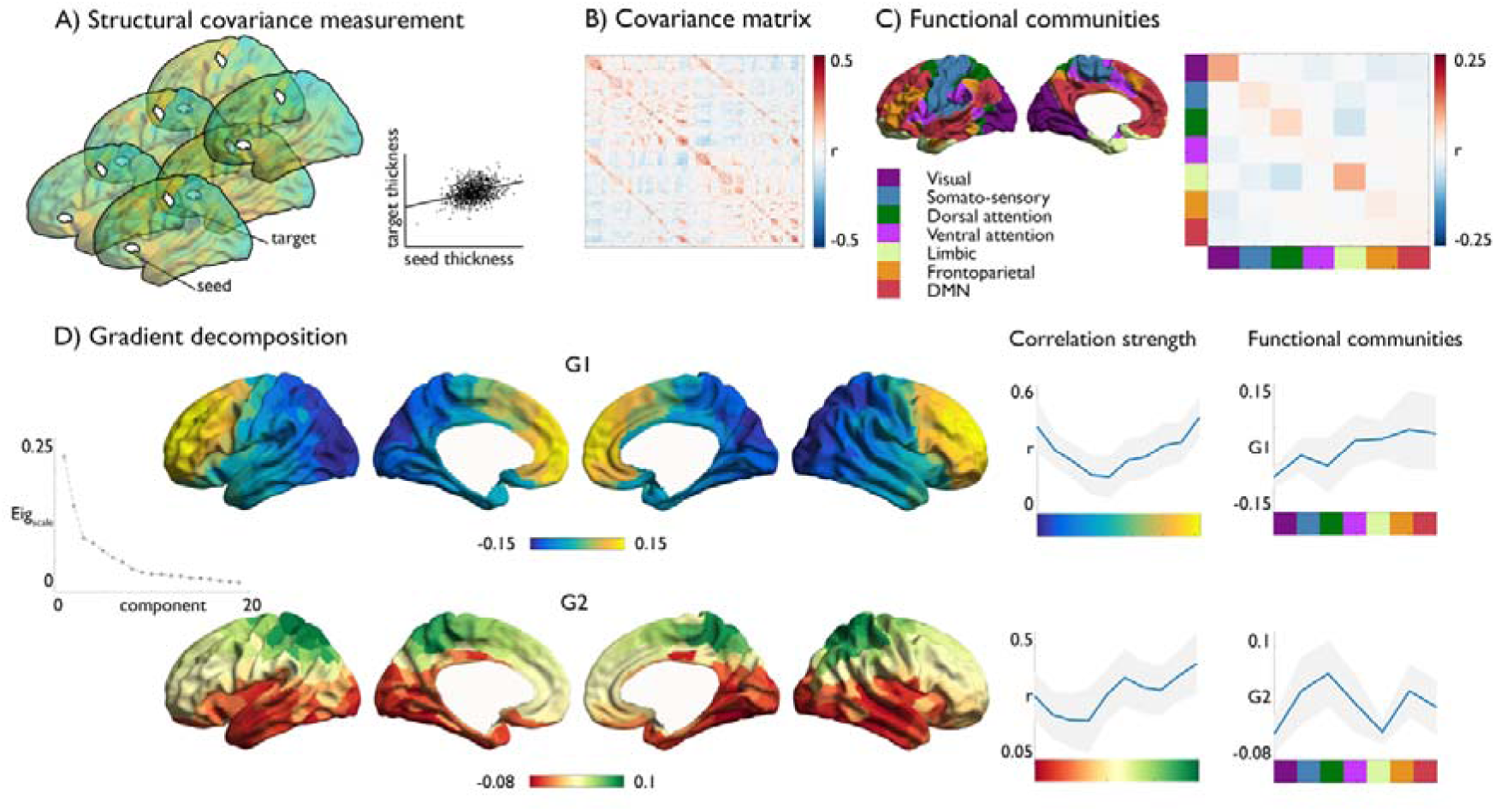
Large scale organization of structural covariance. **A**) Measuring structural covariance of thickness; **B**) Structural covariance matrix; **C**) mean correlation within functional network community ^42^; **D**) Gradient decomposition, primary (G1) and secondary (G2) macro scale gradient, and their average value in mean covariance strength within binned gradient-level, indicating the covariance between regions at similar gradient level, and gradient values as a function of functional community (color nomenclature according to **C**).

We then implemented diffusion map embedding, a method previously used in function connectivity as well as microstructural covariance networks. Diffusion map embedding allows local and long-distance connections to be projected into a common space ^13, 45^. The resulting components are unitless and identify the position of nodes along the respective embedding axis that encodes the dominant differences in nodes’ connectivity patterns. The principal gradient in structural covariance followed a posterior-anterior trajectory from occipital regions to the frontal cortex and accounted for 17% of the variance in the thickness covariance data. Next, we examined the covariance values as a function of the structural gradient. We divided the structural gradient into 10 equally sized bins and plotted the average values of each structural gradient in each bin. We observed that the principal structural covariance gradient followed a U-shaped pattern with both extreme ends of the gradient showing strongest covariance and intermediate zones showing relative low covariance to regions in the same gradient level (Figure 1C). Topology of covariance showed a correspondence to functional organization, with unimodal regions exhibiting lower gradient values relative to networks associated with higher-order processing (default mode network and frontoparietal network) (Figure 1).

The secondary gradient followed an inferior-superior pattern with endpoints in superior parietal lobe and lingual gyrus respectively and explained 13% of the observed variance. Plotting this gradient in each of the 10 bins according to their gradient values indicated that structural covariance increased along the inferior-superior axis, with highest covariance between regions located within superior parietal cortex. Findings were reproducible in a different dataset (eNKI, n=792, age 8-85yrs) (Supplementary Figure 1) and were observed using different preprocessing pipelines of thickness (CIVET and Freesurfer 6.0) (Supplementary Figure 2, **Supplementary Results**) and parcellation methods (Desikan-Killiany^46^, Glasser-atlas^47^, and Schaefer^44^ 800 parcels, Supplementary Figure 3). Notably, age-related effects moderating structural covariance strength also followed posterior-anterior and inferior-superior axes (Supplementary Figure 1**, Supplementary Results**). The primary and secondary gradients, as well as gradients 3 and 4, showed comparable patterning bilaterally, while gradients 5 to 8 showed lateralization effects (Supplementary Figure 4**, Supplementary Results**). Follow up analysis indicated that the gradients of macro scale organization of cortical thickness existed above and beyond geodesic distance constraints, and aligned with previously reported gradients of functional connectivity and microstructural profile covariance (**Supplementary Results**). Conducting a meta-analysis using the Neurosynth database, we observed marked variation of function along both macro scale organizational gradients of thickness (**Supplementary Results**).

### Macro scale organization of cortical thickness is genetically determined

Having established macro scale organizational patterns of cortical thickness, we next computed the genetic correlation between the 400 cortical regions^44^ in the HCP dataset. Genetic correlation is based on the decomposition of structural covariance in genetic and environmental factors using the genetic similarity between individuals to estimate shared additive genetic effects. Overall, 78±5% of the phenotypic correlation could be attributed to genetic factors and we observed high correlation between thickness covariance and genetic correlation of thickness (r=0.61, p<0.0001) and environmental correlation of thickness (r=0.33, p<0.001) across all nodes (Supplementary Figure 5). Patterns of genetic correlation were highest within, rather than between, functional communities (Figure 2A, Supplementary Figure 5, Supplementary Table 2). Though to a lesser extent, this was also the case for environmental influences (Figure 2B, Supplementary Table 3).

**Fig 2.**
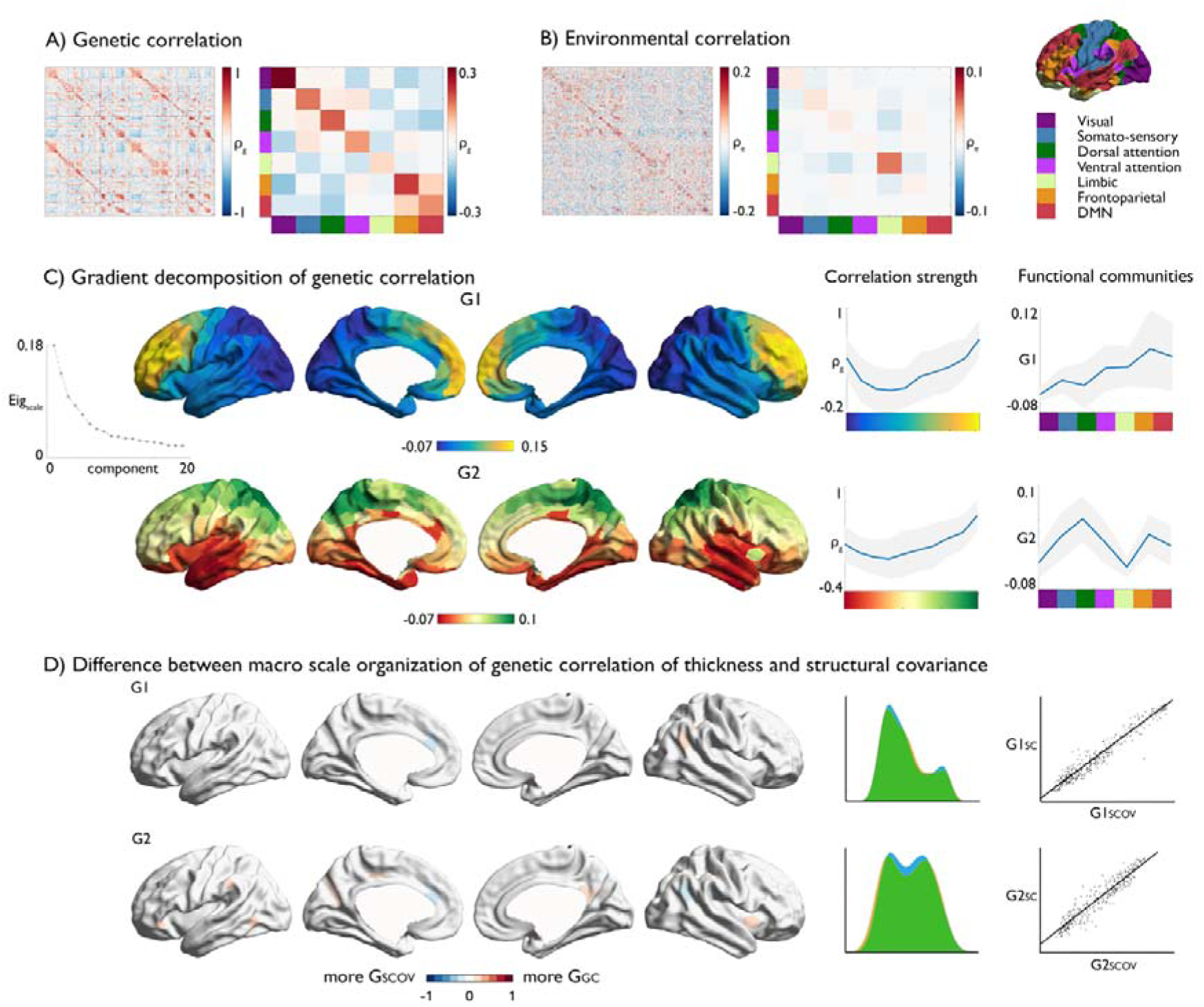
Large scale organization of genetic correlation of cortical thickness. **A**) Genetic correlation of local cortical thickness; i) mean genetic correlation between functional communities ^42^; **B**) Environmental correlation of cortical thickness; i) mean environmental correlation between functional communities ^42^; **C**) Gradient decomposition, primary and secondary macro scale gradient, and their average value in i). mean genetic correlation strength within binned gradient-level; ii). functional communities; **D**). Parcel-wise difference between the structural covariance gradients (G_SCOV_) and the genetic correlation gradients (G_GC_). Blue indicates higher gradient ranking in G_SCOV_, red indicates higher gradient ranking in G_GC_, as well as density plot and scatter of gradient values.

Performing whole-brain gradient decomposition on the genetic correlation maps, we observed almost identical large-scale gradients as in the structural covariance (Structural covariance G1 versus genetic correlation G1: r=0.97, Structural covariance G2 versus genetic correlation G2:r=0.95). The primary genetic gradient explained 18% of the variance, traversing a posterior-anterior axis. Probing the within-gradient genetic correlation, we observed that both end points of the primary gradient showed highest genetic correlation to regions at the same level of the gradient, with the strongest genetic correlation observed in the frontal cortex (Supplementary Figure 6, Supplementary Figure 7). The secondary gradient explained 14% of the variance, and, reflected a similar inferior-superior axis as was seen in the structural covariance gradients. Both organizational axes varied as a function of functional community, suggesting a relationship between the topological organization of genetic correlation of thickness and functional organization. Environmental correlations, explaining 15% of variance of the thickness covariance, were organized along a rostral-caudal and inferior-superior axis as well, explaining 13% and 11% of the variance respectively (Supplementary Figure 8).

### Macro scale organization of cortical thickness in macaques

Thus far our analysis suggests that the macro scale organization of cortical structural covariance in humans shows evidence of high degree of concordance amongst identical twins suggesting a strong genetic influence. Our next analysis evaluated the genetic contribution to macroscale dimensions of cortical structure by examining its phylogenetic stability. To achieve this goal, we examined the topology of large-scale gradients in 41 macaque monkeys from the PRIMatE Data Exchange (PRIME-DE) ^43^. We created a structural covariance matrix based on cortical thickness of 41 macaques, using parcels based on the Markov atlas^48^ and applied a similar analysis as for humans (see Methods). The principal and secondary gradient of the macaque monkey are presented in Figure 3. Similar to the gradients of structural covariance in humans, we observed that the topological organization of macaque monkey’s structural covariance was also well described by both a posterior-anterior and inferior-superior component. In macaques the ordering of the components was reversed with the inferior-superior gradient explained 17% of the variance, whereas the posterior-anterior gradient explained 12% of the variance. The primary gradient stretched from inferior anterior temporal to sensory-motor cortex, and the secondary gradient stretched from sensory-motor to frontal cortex.

**Fig. 3.**
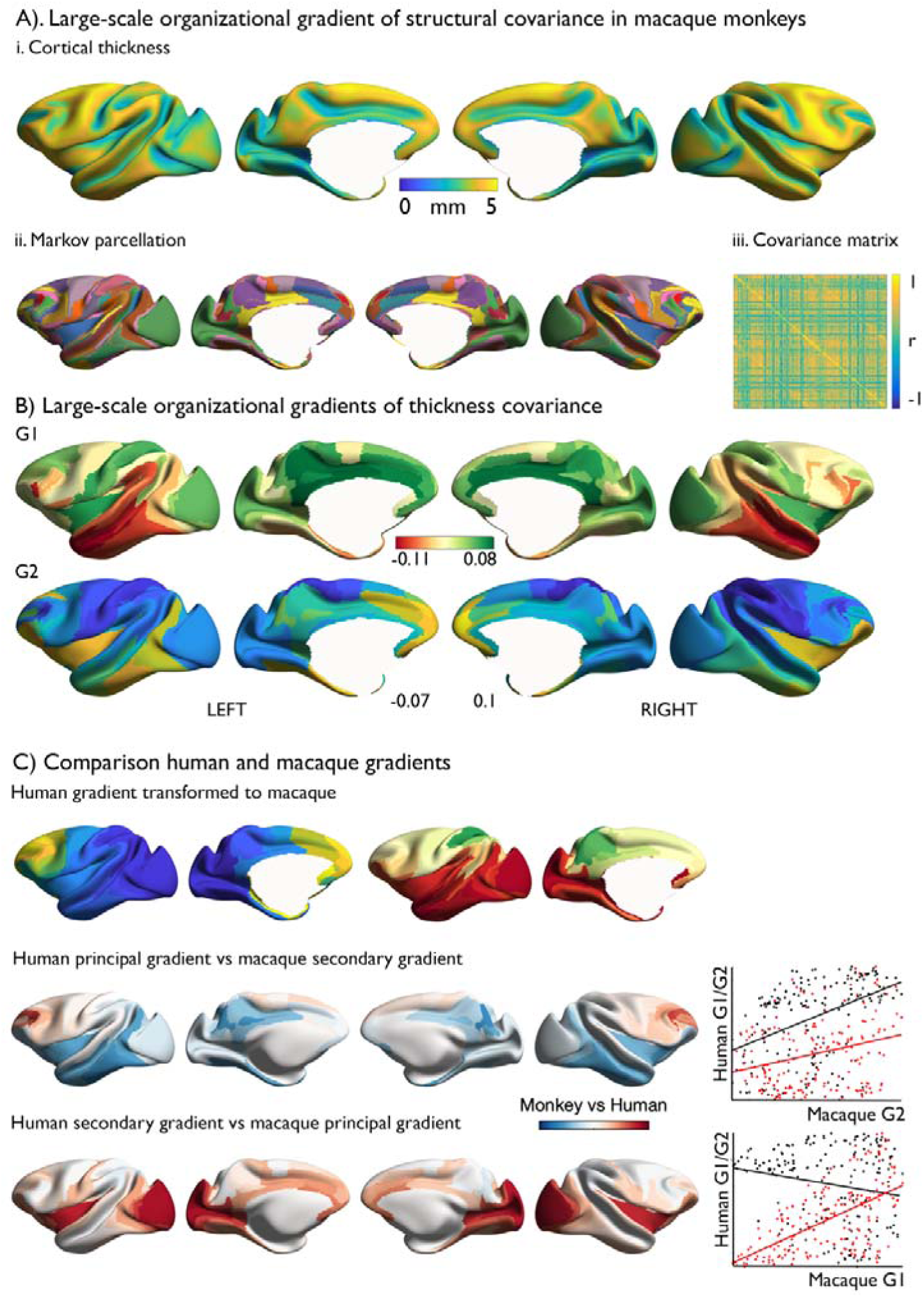
Structural covariance gradient in macaque monkeys. **A**) Mean cortical thickness in 41 macaques from three independent sites (Davis, Oxford, and Newcastle); ii. Markov parcellation^48^; iii. Structural covariance matrix controlling for site. **B)**. Gradient decomposition: primary gradient (G1) and secondary gradient (G2); **C)**. Comparison of human and macaque gradients. Red indicated a higher gradient ranking in humans, whereas blue indicates a higher gradient ranking in macaques. Scatter plots indicate the association between human posterior -anterior covariance gradient (G1, black) and human inferior- superior covariance (G2, red) and macaque principal gradient (G1, upper scatterplot) and secondary gradient (G2, lower scatterplot).

Last, using an innovative approach to perform cross species alignment (weighted functional-alignment) ^49^ we transformed human gradients to macaque cortex and compared them with the gradients in macaques directly. We observed strong similarity between the posterior-anterior gradient (r=0.52, [0.41, 0.61], p<0.0001) and inferior-superior gradients in humans and macaques (r=0.60, [0.49, 0.70]), p<0.0001). Notably, these similarities were stronger than between posterior-anterior gradient in humans and inferior-superior gradient in macaques (r=-0.08, [-0.23, 0.04], p=ns) or inferior-superior gradient in humans and posterior-anterior gradient in macaques (r=0.24, [0.08, 0.37], p=0.001).

### Macro scale organization of cortical thickness and the theory of dual origin

Finally, we studied the genetic ontogeny of macro scale organization of cortical thickness in light of the dual origin theory of cortical development. This perspective assumes that cortical areas develop from waves of laminar differentiation that have their origin in either the piriform cortex (paleo-cortex) or the hippocampus (archi-cortex). The theory was established on histological investigations of the adult cortex of various reptiles and mammals ^3, 17–20, 50^. We evaluated the previously reported gradients in humans and macaques with respect to the geodesic distance from the paleo-cortex (olfactory cortex) and the archi-cortex (hippocampus) (similar to previous work ^16^).

In humans, the paleocortex was defined by the paleocortex, and the archi-cortex was defined by hippocampus, pre-subiculum, area 33’, and retrosplenial complex. We computed the average geodesic distance from these ROIs (Figure 4a) and evaluated its association to the principal and secondary gradient of genetic correlation of thickness (based on Figure 2). We observed a dissociation between distance from paleo-cortex in inferior and superior proportions of the inferior-superior gradient (statistical energy-test^51^: p<0.001). And, using spin-tests to account for spatial autocorrelation ^52^, we observed a negative relation between the paleo-cortex distance map and inferior-superior gradient level (r_spin_=-0.78, p<0.01), suggesting that the macro scale structural organization varies gradually as a function of paleo-cortex distance. Contrarily, there was positive relationship between inferior and superior proportions of the inferior-superior gradient and archi-cortex distance (energy-test: p<0.003) and a negative, but non-significant, linear relationship between this gradient and archi-cortex distance (r_spin_ =-0.24, p>0.1). We did not observe a consistent association between the dual origin and the posterior-anterior gradient (archi-cortex distance: energy-test: p>0.1, r_spin_ =0.12, p>0.1; paleo-cortex: energy-test: p<0.0001, r_spin_ =-0.43, p>0.1). Evaluating genetic correlation as a function of paleo- and archi-cortex distance, we observed that genetic correlation varied as a function of distance from both origins.

**Fig. 4.**
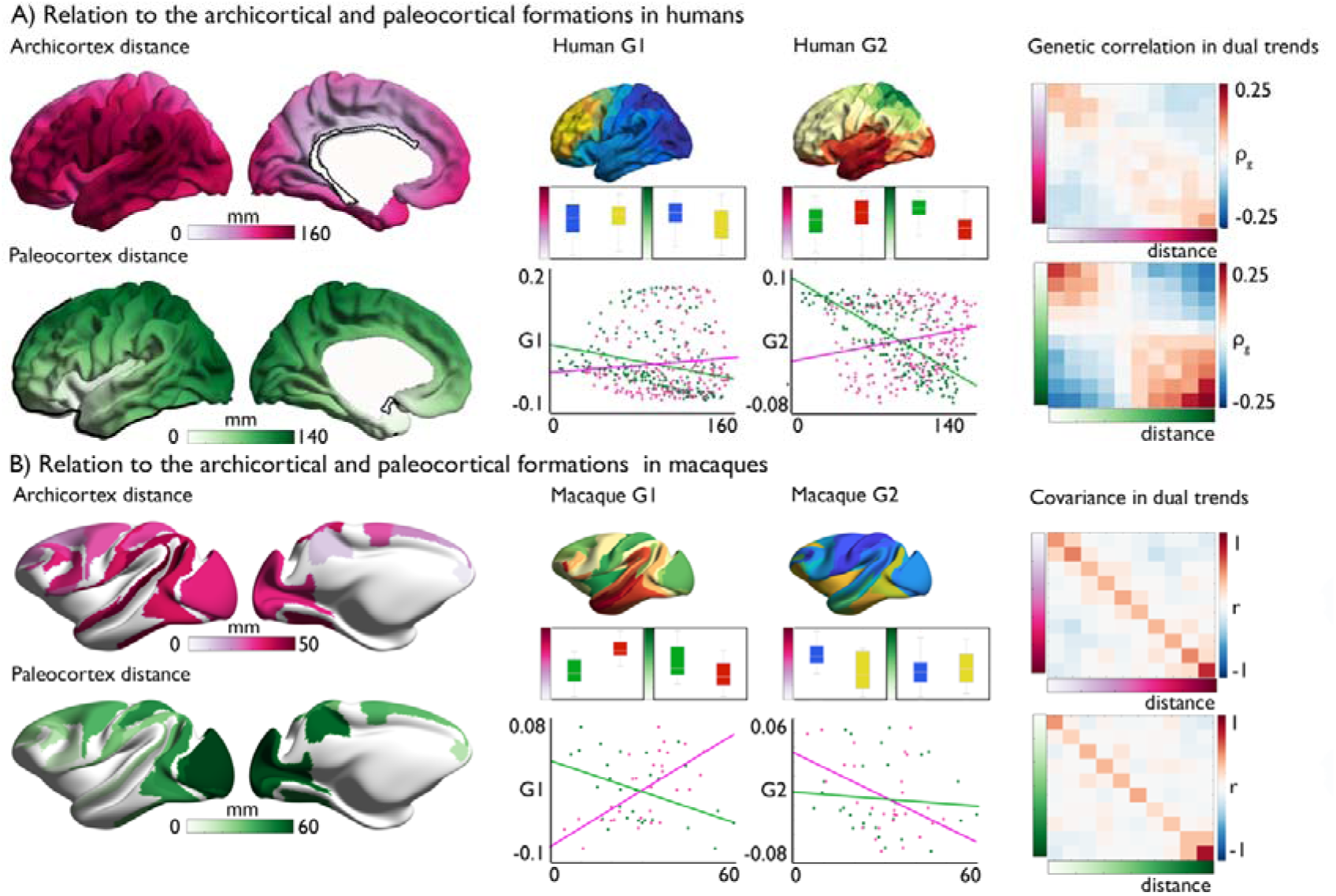
Cross-species topology of covariance as a function of the dual origin theory. **A)**. Left: distance from archi-cortex and paleo-cortex in humans; Middle: Association between G1 and G2 of genetic correlation of thickness and distance from archi-cortex and paleo-cortex in humans (two binned gradients, as well as linear relationship); Right: genetic correlation as a function of archi- and paleo-cortex distance; **B)**. Left: Distance from archi- cortex and paleo-cortex in macaque monkeys ^16^; Middle: Association between G1 and G2 of thickness covariance and distance from archi-cortex and paleo-cortex in macaque monkeys (two binned gradients, as well as linear relationship); Right: structural covariance as a function of archi- and paleo-cortex distance^16^.

We performed a similar analysis in macaque monkeys, using the distance from archi- and paleo-cortex reported by Goulas et al.^16^. We observed that the inferior-superior gradient in structural covariance showed a positive association with archi-cortex distance (energy-test: p<0.002, r=0.64, p<0.0001) and a negative association with paleocortex distance (energy-test: p=ns, r=-0.40, p<0.04). Again, we did not observe a consistent association between the dual origin and the posterior-anterior gradient (archi-cortex distance: energy-test: p<0.02, r=-0.31, p>0.1; paleo-cortex distance: energy-test: p>0.1, r =-0.14, p>0.1).

## Discussion

The cortical mantle is organized along axes that reflect systematic variations in brain structure and function such as laminar differentiation, gene expression, structural and functional connectivity. Although the importance of macro scale axes of cortical organization for human cognition and disorder are now recognized, the degree to which these topological features of the cerebral cortex are genetically determined remains incompletely understood. Our current study provided converging evidence that genetic influences contribute to the spatial organization of macro scale structural features of the cortex. In humans we found two robust topological patterns of macro scale organization of thickness; a posterior-anterior and an inferior-superior gradient, and almost identical organization patterns were observed when assessing genetic correlation of thickness. Furthermore, we found that similar patterns of macro scale organization of cortical thickness as are seen in humans were present in macaque monkeys. Last, we show that both in humans and macaques the inferior-superior axis could be aligned with organization patterns expected based on the theory of dual origin, providing a neurogenetic basis for observed topological patterns. Together, these different analyses provide converging evidence of the important role that genetic influences play in determining the macro scale organization of the cortex.

Our study builds on a growing body of evidence describing the organizational axes that determine the macro scale organization of specific brain features such as myeloarchitecture ^2–6^, cytoarchitecture ^7–10^, laminar origin of connections ^11, 12^ ^10^, functional connectivity ^13^, cortical thickness ^14^, and gene expression ^6, 15^. Together these studies indicate that the transition from cortical areas with less to more laminar differentiation constitutes major axis of cortical organization across which cortical features systematically vary ^6, 12, 13^. These variations have functional and behavioural ramifications ^1, 13^ and the systematic topological organization of the cerebral cortex has been proposed to reflect an architecture which optimize the balance of externally and internally oriented functioning, which is critical for flexibility of human cognition^13, 22, 26^. In the current study we uncovered two major topological axes in macro scale organization of thickness, of which the posterior-anterior gradient explained the greatest amount of variance in humans. Various studies ^53–56^ have demonstrated a posterior-anterior gradient in neuron number in the cortex of a broad range of mammalian species, including rodents, marsupials, and non-human primates ^1, 54, 57^. Neuron numbers are high in posterior portions of the cortex, such as the occipital lobe, and gradually decreases toward more anterior regions. The difference in neuronal numbers has been found to relate to the temporal sequence of neurogenesis ^55, 57^, whereas posterior regions undergo a high number of cell cycles, which accounts for the higher number of neurons in these areas, in anterior regions more time is devoted to the growth of large neurons with many connections ^58^. The posterior-anterior gradient therefore might signify a shift in computational capacity, from a high number of processing units in caudal regions, to a lower number of highly connected units in rostral regions ^55^. Functionally, human imaging studies have placed representation of stimulus properties posteriorly, involving local computations, and more complex operations, involving integration of various functions, anteriorly ^59–61^.

A second organizational axis in macro scale organization of thickness was identified that followed an inferior-superior pattern in humans and macaque monkeys. Inferior-superior (dorso-ventral) patterning is a key organizational principle during embryonic development of the central nervous system ^3, 16–20, 62, 63^ and dorsal-ventral dichotomies have been reported in macaques ^9 64 65^ and humans ^66^. Notably, the inferior-superior axis differentially related to distance from paleo- and archi-cortex respectively, aligning the inferior-superior axis in macro scale organization of thickness with the dual origin theory. This convergence suggests that our method captures at a macro scale how regions, which could be reasonably distant in space can be affiliated because they share similar origins ^9, 16, 20^. The emergence of the dual connectional trends might be rooted in two patterns centers in the developing pallium, resulting in two opposing neurogenetic gradients ^2^. Both ventral and dorsal systems have been proposed to relate to differentiable functional processes. Whereas the dorsal system has been proposed to relate to time, space, and motility, the ventral system has been associated with assigning meaning and motivation ^66–68^.

We observed differential ordering of posterior-anterior and inferior-superior gradients in humans and macaques. Whereas in humans the principal gradient traversed a posterior-anterior trajectory, we observed that in macaques this gradient was only the second description of shared variance. This difference might reflect the difference in the timing of cortical expansion between humans and macaques. For example, it has been shown that in the macaque monkey, neurogenesis ends about 20 days earlier in the rostral pole than in the most caudal regions ^69^, in humans, however, a posterior-anterior difference of up to 70 days has been predicted ^57^. It is possible that difference in timing of neurogenesis might describe why the same axis of organization can be more or less pronounced in different species. Previous work, using the same sample of macaques, has shown that similarity in functional cortical organization between humans and macaques decreases with geodesic distance from unimodal systems, culminates in the greater differences in posterior regions of the default network. It is possible this functional difference emerges from the different balance of the structural organizational patterns between macaques and humans. Notably, it has been suggested that the evolution of the globular shape of the human brain is related to genes involved in neurogenesis and myelination ^70^, resulting in relatively globular shape of the brain in modern humans relative to its ancestors. It will be important for future work to explore whether differences in the emphasis placed on similar organizational patterns across different species can describe the evolutionary differences in cognitive functions between humans and other primates.

Follow up analysis indicated the posterior-anterior and inferior-superior organization gradients in macro scale organization of thickness is similar to previously described gradients in microstructural profile covariance ^6^ and functional connectivity ^13^. The posterior-anterior gradient related to T1wT2w contrast in all layers. This is in line with previous in vivo and post-mortem evidence of an increase of mean myelin from polar towards sensory regions ^71, 72^. The dorsal-ventral dissociation was only observed in the upper two strata, with ventral regions relating to lower T1wT2 contrast than dorsal regions. Difference in upper and lower strata T1wT2w contrast has been summarized using “skewness”, indicating regions with high difference between upper and lower layers would have a low skewness, whereas regions with a small difference between upper and lower layers having a high skewness ^73^. Dorsal regions including the sensory-motor cortex have been reported to have a low skewness, indicating a high difference in myelin between upper and lower layers. It is possible that the dorsal-ventral patterning of myelin in the upper layers reflects a dissociation in information processing, with sensory agranular regions providing feedforward information and project locally, whereas ventral, more granular paralimbic, regions are involved in feedback processing and project from infragranular layers ^74, 75^. Additionally, we found comparable topologies in microstructural profile covariance and macro scale organization of thickness, in line with previous evidence that thickness topology relates to microstructural differentiation ^14, 76^. Notably, both posterior-anterior macro scale organization patterns, as well as the combination of both the posterior-anterior and inferior-superior gradient showed a positive relation primary organizational axis of functional connectivity at rest. Our observation that a combination of gradients associated with differing neurogenetic and developmental mechanisms puts forward the hypothesis that functional organization arises through the combination of multiple structural organizational axes, and, as such, creating an architecture which optimize the balance of externally and internally oriented functioning, which is critical for flexibility of human cognition.

Understanding of large-scale organization of brain structure may offer a novel and compelling model to evaluate to impact and progression of pathology. For example, it has been suggested that Parkinson’s and Alzheimer’s disease follows a staging trajectory, with different regions and networks affected at different stages of the disorder ^27, 28^, and its sequence determined by underlying anatomical axes. Parkinson’s is assumed to show early disruptions in the lower brain stem, followed later disruption in other midbrain structures, meso-cortex and allocortex. Final stages of the disorder are characterized by disruptions in sensory-motor areas. We note that this sequence of deficits is similar to the inferior-superior axis, suggesting that understanding this feature of cortical organization may also help understand the apparent sequence of deficits in Parkinson’s disease. Future work should therefore consider whether the macro scale patterns of that our analysis shows reflect the contribution of genetic influences may shed light on specific orderly sequences in symptoms that underpins Parkinson’s disease, as well as other neurodegenerative conditions.

To conclude, our novel results establish that two major organizational axes in macro scale organization of thickness in human and non-human primates that are likely to be at least partially influenced by genes. We found a principal gradient stretched from posterior to anterior cortical areas, whereas a secondary gradient traversed along an inferior-superior gradient, and aligned with theories on the dual origin of the cortex. Combined, our observations provide direct evidence of a genetic basis of macro scale organizational patterns. It is of note that our findings were made possible thanks to open data initiatives. These initiatives offer the neuroimaging and network neuroscience communities an unprecedented access to large datasets for the investigation of human and non-human brains and for the cross-validation of observations across data-sets and methods. Uncovering the organizational axis of the human cerebral cortex provides insights in the neurogenetic processes shaping its structural and functional organization and its relation to human cognition. Such axes can be utilized to evaluate disease progression as well as disseminate potential neurogenetic origins of abnormal cortical development.

## Materials and methods

### HCP sample

#### Participants and study design

For our analysis we used the publicly available data from the Human Connectome Project S900 release (HCP; http://www.humanconnectome.org/), which comprised data from 970 individuals (542 females), 226 MZ twins, 147 DZ twins, and 597 singletons, with mean age 28.8 years (SD = 3.7, range = 22–37). We included individuals for whom the scans and data had been released (humanconnectome.org) after passing the HCP quality control and assurance standards ^77^. The full set of inclusion and exclusion criteria are described elsewhere ^78, 79^. In short, the primary participant pool comes from healthy individuals born in Missouri to families that include twins, based on data from the Missouri Department of Health and Senior Services Bureau of Vital Records. Additional recruiting efforts were used to ensure participants broadly reflect ethnic and racial composition of the U.S. population. Healthy is broadly defined, in order to gain a sample generally representative of the population at large. Sibships with individuals having severe neurodevelopmental disorders (e.g., autism), documented neuropsychiatric disorders (e.g. schizophrenia or depression) or neurologic disorders (e.g. Parkinson’s disease) are excluded, as well as individuals with diabetes or high blood pressure. Twins born prior 34 weeks of gestation and non-twins born prior 37 weeks of gestation are excluded as well. After removing individuals with missing structural imaging data our sample consisted of 899 (504 females) individuals (including 220 MZ-twins and 135 DZ-twins) with a mean age of 28.8 years (SD =3.7, range =22-37).

#### Structural imaging processing

MRI protocols of the HCP are previously described in^78, 79^. In short, MRI data used in the study were acquired on the HCP’s custom 3T Siemens Skyra equipped with a 32-channel head coil. Two T1w images with identical parameters were acquired using a 3D-MPRAGE sequence (0.7 mm isotropic voxels, matrix = 320 × 320, 256 sagittal slices; TR = 2,400 ms, TE = 2.14 ms, TI = 1,000 ms, flip angle = 8°; iPAT = 2). Two T2w images were acquired using a 3D T2-SPACE sequence with identical geometry (TR = 3,200 ms, TE = 565 ms, variable flip angle; iPAT = 2). T1w and T2w scans were acquired on the same day. The pipeline used to obtain the Freesurfer-segmentation is described in detail in a previous article ^78^ and is recommended for the HCP-data. The pre-processing steps included co-registration of T1- and T2-weighted scans, B1 (bias field) correction, and segmentation and surface reconstruction using FreeSurfer version 5.3-HCP to estimate cortical thickness.

In addition to assess robustness and replicability of the results across different surface estimation pipelines, cortical thickness estimates were further estimated using FreeSurfer version 6.0 and CIVET. For both these additional analyses, only bias-corrected T1-weighted data were used as the input. FreeSurfer version 6.0 was performed using the default recon-all options. Surface-extraction and cortical thickness estimation using CIVET were performed using version 2.1.1 (http://www.bic.mni.mcgill.ca/ServicesSoftware/CIVET). The non-uniformity artefacts were corrected with the N3 algorithm (Sled et al., 1998) using the recommended N3 spline distance of 125mm for 3T T1-weighted scans. Cortical thickness was then measured as the distance between the estimated “white” and “grey” cortical surfaces, in the native space framework of the original MR images, using the same approach that is used in FreeSurfer^80^.

#### Parcellation approach

We used a parcellation scheme^44^ based on the combination of a local gradient approach and a global similarity approach using a gradient-weighted Markov Random models. The parcellation has been extensively evaluated with regards to stability and convergence with histological mapping and alternative parcellations. In the context of the current study, we focus on the granularity of 400 parcels, as averaging will improve signal-to-noise. In order to improve signal-to-noise and improve analysis speed, we opted to average unsmoothed structural data within each parcel. Thus, cortical thickness of each ROI was estimated as the trimmed mean (10 percent trim). Findings were additionally evaluated using different parcellation schemes using the 800 parcel Schaefer^44^ solution, as well as the Glasser atlas^47^ based on myelo-architecture and the Desikan-Killiany^46^ atlas.

#### Gradient decomposition

In line with previous studies ^5, 13^ the structural covariance and genetic correlation matrix, as well as age-related t-maps, were proportionally thresholded at 90% per row and converted into a normalized angle matrix using the BrainSpace toolbox for matlab ^52^. Diffusion map embedding^45^, a non-linear manifold learning technique, identified principal gradient components, explaining structural covariance variance in descending order (each of 1 × 400). In brief, the algorithm estimates a low-dimensional embedding from a high-dimensional affinity matrix. In this space, cortical nodes that are strongly interconnected by either many supra-threshold edges or few very strong edges are closer together, whereas nodes with little or no covariance are farther apart. The name of this approach, which belongs to the family of graph Laplacians, derives from the equivalence of the Euclidean distance between points in the embedded space and the diffusion distance between probability distributions centered at those points. It is controlled by a single parameter α, which controls the influence of the density of sampling points on the manifold (α = 0, maximal influence; α = 1, no influence). Based on previous work ^5, 13^ we followed recommendations and set α = 0.5, a choice that α retains the global relations between data points in the embedded space and has been suggested to be relatively robust to noise in the covariance matrix. Gradients were mapped onto fsaverage surface visualized using SurfStat (http://mica-mni.github.io/surfstat)^77^ and we assessed the amount of variance explained. To show how the principal and secondary gradient of covariance/genetic correlation relates to systematic variations in functional organization ^42^, we calculated and plotted the mean covariance profiles within ten equally sized discrete bins of the respective gradient. To evaluate correlation between macrostructural gradients we used spin permutations ^35^.

#### Genetic correlation analysis

To investigate the genetic correlation of brain structure, we analyzed 400 parcels of cortical thickness in a twin-based genetic correlation analysis. As in previous studies ^81^, the quantitative genetic analyses were conducted using Sequential Oligogenic Linkage Analysis Routines (SOLAR) ^82^. SOLAR uses maximum likelihood variance-decomposition methods to determine the relative importance of familial and environmental influences on a phenotype by modeling the covariance among family members as a function of genetic proximity. This approach can handle pedigrees of arbitrary size and complexity and thus, is optimally efficient with regard to extracting maximal genetic information. To ensure that our cortical thickness parcels were conform to the assumptions of normality, an inverse normal transformation was applied^81^.

Heritability (*h*^2^) represents the portion of the phenotypic variance (σ^2^_p_) accounted for by the total additive genetic variance (σ^2^_g_), i.e., *h*^2^ = σ^2^_g_/σ^2^_p_. Phenotypes exhibiting stronger covariances between genetically more similar individuals than between genetically less similar individuals have higher heritability. Within SOLAR, this is assessed by contrasting the observed covariance matrices for a neuroimaging measure with the structure of the covariance matrix predicted by kinship. Heritability analyses were conducted with simultaneous estimation for the effects of potential covariates. For this study, we included covariates including global thickness, age, and sex.

To determine if shared variations in cortical thickness were influenced by the same genetic factors, genetic correlation analyses were conducted. More formally, bivariate polygenic analyses were performed to estimate genetic (ρ_g_) and environmental (ρ_e_) correlations, based on the phenotypic correlation (ρ_p_), between brain structure and personality with the following formula: 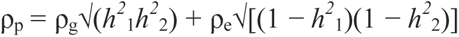 here where *h_1_^2^* and *h_2_^2^* are the heritability of the parcel-based cortical thickness. The significance of these correlations was tested by comparing the log likelihood for two restricted models (with either ρ_g_ or ρ_e_ constrained to be equal to 0) against the log likelihood for the model in which these parameters were estimated. A significant genetic correlation (corrected for multiple comparisons using Bonferroni correction) is evidence suggesting that (a proportion of) both phenotypes are influenced by a gene or set of genes^83^. To compute the contribution of genetic effects relative to the phenotypic correlation, we computed the contribution of the genetic path to the phenotypic correlation 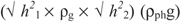 divided by the phenotypic correlation. For the relative contribution of environmental correlation to the phenotypic correlation we computed 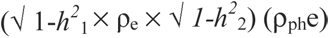 divided by the phenotypic correlation^84^.

#### Geodesic distance

Geodesic distance was computed between each vertex in fsaverge5 space using the Eucledian coordinates of the vertices, creating a 20484 x 20484 distance matrix. Only ipsilateral distance was considered. Following distances between parcels were computed by taking the average distance between both parcels. We evaluated the macro scale organization of thickness while controlling for distance by multiplying the covariance strength by the distance between the respective parcels.

#### Comparisons between gradients and modalities

To make comparisons across gradient and distance maps, we used spin-tests to control for spatial autocorrelation when possible^85^. Difference between the two distributions of archi- and paleo-cortex distance and macro scale organizational gradients were assessed using statistical energy test, a non-parametric statistic for two sample comparisons ^51^ (https://github.com/brian-lau/multdist/blob/master/minentest.m) and statistical significance was assessed with permutation tests (1000).

#### Macaque sample

We used the MRI data from the recently formed NHP data sharing consortium PRIME-DE [http://fcon_1000.projects.nitrc.org/indi/indiPRIME.html]. Three cohorts of macaque monkeys were included in the present study (Newcastle University, Oxford University, and University of California, Davis).

##### Oxford data

The full data set consisted of 20 rhesus macaque monkeys (*macaca mulatta*) scanned on a 3T scanner with 4-channel coil. The data were collected while the animals were under anesthesia. Briefly, the macaque was sedated with intramuscular injection of ketamine (10 mg/kg) combined with either xylazine (0.125-0.25 mg/kg) or midazolam (0.1mg/kg) and buprenorphine (0.01 mg/kg). Additionally, macaques received injections of atropine (0.05 mg/kg, i.m.), meloxicam (0.2 mg/kg, i.v.), and ranitidine (0.05 mg/kg, i.v.). The anesthesia was maintained with isoflurane. The details of the scan and anesthesia procedures were described in ^86^ and the PRIME DE website (http://fcon_1000.projects.nitrc.org/indi/PRIME/oxford.html).

##### UC-Davis Data

The full data set consisted of 19 rhesus macaque monkeys (macaca mulatta, all female, age=20.38 ± 0.93 years, weight=9.70 ± 1.58 kg) scanned on a Siemens Skyra 3T with 4-channel clamshell coil. All the animals were scanned under anesthesia. In brief, the macaques were sedated with injection of ketamine (10 mg/kg), dexmedetomidine (0.01 mg/kg), and buprenorphine (0.01 mg/kg). The anesthesia was maintained with isoflurane at 1-2%. The details of the scan and anesthesia protocol can be found at (http://fcon_1000.projects.nitrc.org/indi/PRIME/ucdavis.html).

##### Newcastle data

The full data set consisted of 14 rhesus macaque monkeys (*macaca mulatta*) scanned on a Vertical Bruker 4.7T primate dedicated scanner. We restricted our analysis to 10 animals (8 males, age=8.28±2.33, weight=11.76±3.38) for whom two awake resting-state fMRI scans were required. The structural T1-weighted images were acquired using MDEFT sequence with 0.6×0.6×0.6mm resolution, TE=6ms, TR=750ms.

##### MRI data processing

The structural processing includes 1) spatial denoising by a non-local mean filtering operation ^87^, 2) brain extraction using ANTs registration with a reference brain mask followed by manually editing to fix the incorrect volume (ITK-SNAP, www.itksnap.org) ^88^; 3) tissue segmentation and surface reconstruction (FreeSurfer)^89, 90^; 4) the native white matter and pial surfaces were registered to the Yerkes19 macaque surface template ^91^.

#### Quality control

We excluded macaque monkeys that showed a hemispheric difference of more 0.2 cm (UC Davis (0); Oxford (7), Newcastle (5)) for our final analysis, as gradient models were estimated based on covariance of ipsi- and contra-lateral covariance.

#### Gradient analysis

First we constructed a covariance matrix, controlling for dataset site and global thickness. Following we performed gradient analysis analogue to described in humans. *Alignment of human gradients to macaque gradients:* To evaluate the similarity between human and macaque gradients we transformed the human gradient to macaque cortex based on a functional-alignment techniques recently developed. This method leverages advances in representing functional organization in high-dimensional common space and provides a transformation between human and macaque cortices. ^49^.

*Archi-paleo cortex distance:* Distance from the archi – and paleo cortex was computed in Goulas et al., 2019 ^16^.

#### Cortical microstructure and microstructural covariance networks

We estimated MPC using myelin-sensitive MRI (MPCMRI), in line with the previously reported protocol ^5^, in our main sample (HCP S900). The myelin-sensitive contrast was T1w/T2w from the HCP minimal processing pipeline, which uses the T2w to correct for inhomogeneities in the T1w image. We generated 12 equivolumetric surfaces between the outer and inner cortical surfaces ^92^. The equivolumetric model compensates for cortical folding by varying the Euclidean distance ρ between pairs of intracortical surfaces throughout the cortex to preserve the fractional volume between surfaces ^93^. ρ was calculated as follows for each surface (1):

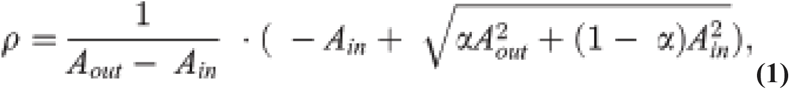

in which α represents a fraction of the total volume of the segment accounted for by the surface, while A_out_ and A_in_ represents the surface area of the outer and inner cortical surfaces, respectively. We systematically sampled T1w/T2w values along 64,984 linked vertices from the outer to the inner surface across the whole cortex. Following we computed the average value of T1w/T2 in each of the 400 parcels of the Schaefer atlas^44^. In turn, MPC_MRI_*(i*,*j)* for a given pair of parcels *i* and *j* is defined by (5):

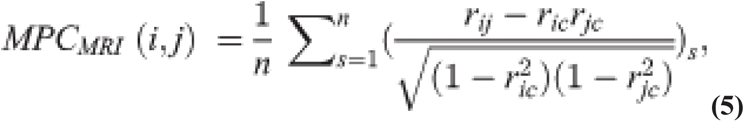

in which *s* is a participant and *n* is the number of participants. We used the MPC_MRI_ to (re-) compute the gradient of microstructure.

#### Functional connectivity gradient

The functional connectivity gradient was downloaded from (https://www.neuroconnlab.org) computed as part of ^13^, based on 820 individuals from the HCP S900 release. As the gradient was reported at the fs_32k standard space surface, values were resampled for the Schaefer 400 parcellation for further analysis.

### Replication sample<u>: eNKI</u>

#### Participants and study design

To evaluate the cross-sample reproducibility of observations we additionally investigated correspondence between personality and cortical brain structure in the enhanced Nathan Kline Institute-Rockland Sample (NKI). The sample was made available by the Nathan-Kline Institute (NKY, NY, USA), as part of the ‘*enhanced NKI-Rockland sample*’ (https://www.ncbi.nlm.nih.gov/pmc/articles/PMC3472598/). In short, eNKI was designed to yield a community-ascertained, lifespan sample in which age, ethnicity, and socioeconomic status are representative of Rockland County, New York, U.S.A. ZIP-code based recruitment and enrollments efforts were being used to avoid over-representation of any portion of the community. Participants below 6 years were excluded to balance data losses with scientific yield, as well as participants above the age of 85, as chronic illness was observed to dramatically increase after this age. All approvals regarding human subjects’ studies were sought following NKI procedures. Scans were acquired from the International Neuroimaging Data Sharing Initiative (INDI) online database http://fcon_1000.projects.nitrc.org/indi/enhanced/studies.html For our phenotypic analyses, we selected individuals with complete personality and imaging data. Our sample for phenotypic correlations consisted of 799 (400 females) individuals with a mean age of 41.1 years (SD =20.3, range =12-85).

#### Structural imaging processing

3D magnetization-prepared rapid gradient-echo imaging (3D MP-RAGE) structural scans^88^ were acquired using a 3.0-T Siemens Trio scanner with TR=2500 ms, TE=3.5-ms, Hz/Px, field of view=256 × 256-mm, flip angle=8°, voxel size=1.0 × 1.0 × 1.0-mm. More details on image acquisition are available at http://fcon_1000.projects.nitrc.org/indi/enhanced/studies.html. All T1 scans were pre-processed using the Freesurfer software library (http://surfer.nmr.mgh.harvard.edu/) version 6.0.0 ^80, 89, 90, 94^ to compute cortical thickness. Next, the individual cortical thickness and surface area maps were standardized to fsaverage5 for further analysis. Segmentations were visually inspected for anatomical errors (S.L.V.).

#### Modulation of structural covariance of thickness by age

In the eNKI sample, we also computed the modulation of structural covariance by probing the interaction of covariance by age in the following model:

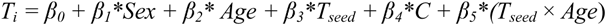

Following the parcel to parcel t-maps were used to compute large-scale gradients age-related changes in covariance.

#### Replication: cortical thickness methodology

Cortical thickness of the individuals of the HCP S1200 release were computed as part of an independent study (Kharabian, under review) and resampled to Schaefer 400 parcels. We utilized the extracted thickness values of FreeSurfer 6.0 to evaluate the stability of observed covariance organization as a function of cortical thickness estimation method. For the FreeSurfer 6.0. analysis of the T1-weighted images in the HCP dataset we used the default recon-all options (version (v) 6.0; (www.surfer.nmr.mgh.harvard.edu)). Moreover, cortical thickness estimation using CIVET were performed using version 2.1.1 (http://www.bic.mni.mcgill.ca/ServicesSoftware/CIVET).

## Data availability

All human data analyzed in this manuscript were obtained from the open-access HCP young adult sample (HCP; http://www.humanconnectome.org/)^79^ and enhanced NKI-Rockland sample (https://www.ncbi.nlm.nih.gov/pmc/articles/PMC3472598/) ^95^. Scans were acquired from the International Neuroimaging Data Sharing Initiative (INDI) online database http://fcon_1000.projects.nitrc.org/indi/enhanced/studies.html. The raw data may not be shared by third parties due to ethics requirements, but can be downloaded directly via the above weblinks. Macaque data was obtained from the recently formed NHP data sharing consortium PRIME-DE [http://fcon_1000.projects.nitrc.org/indi/indiPRIME.html]. Three cohorts of macaque monkeys were included in the present study (Newcastle University, Oxford University, and University of California, Davis). Genetic analyses were performed using Solar Eclipse 8.4.0 (http://www.solar-eclipse-genetics.org), and data on the KING pedigree analysis is available here: https://www.nitrc.org/projects/se_linux/ ^82, 96^. Gradient mapping analyses was based on open-access tools (Brainmap, https://brainspace.readthedocs.io/en/latest/). Surface-wide statistical comparisons and visualizations were carried out using SurfStat https://github.com/MICA-MNI/micaopen/tree/master/surfstat) in combination with colorbrewer (https://github.com/scottclowe/cbrewer2). Both structural covariance and genetic correlation gradients are available at (https://github.com/sofievalk/projects/tree/master/Structure_of_Structure).

## Acknowledgements

We would like to thank the various contributors to the open access databases that our data was downloaded from. Specifically; HCP data were provided by the Human Connectome Project, Washington University, the University of Minnesota, and Oxford University Consortium (Principal Investigators: David Van Essen and Kamil Ugurbil;1U54MH091657) funded by the 16 NIH Institutes and Centers that support the NIH Blueprint for Neuroscience Research; and by the McDonnell Center for Systems Neuroscience at Washington University. For enhanced NKI, we would like to thank the principal support for the enhanced NKI-RS project is provided by the NIMH BRAINS R01MH094639-01 (PI Milham). Funding for key personnel was also provided in part by the New York State Office of Mental Health and Research Foundation for Mental Hygiene. Funding for the decompression and augmentation of administrative and phenotypic protocols provided by a grant from the Child Mind Institute (1FDN2012-1). Additional personnel support provided by the Center for the Developing Brain at the Child Mind Institute, as well as NIMH R01MH081218, R01MH083246, and R21MH084126. Project support also provided by the NKI Center for Advanced Brain Imaging (CABI), the Brain Research Foundation (Chicago, IL), and the Stavros Niarchos Foundation. This study was supported by the Deutsche Forschungsgemeinschaft (DFG, EI 816/21-1), the National Institute of Mental Health (R01-MH074457), the Helmholtz Portfolio Theme “Supercomputing and Modeling for the Human Brain” and the European Union’s Horizon 2020 Research and Innovation Program under Grant Agreement No. 785907 (HBP SGA2). BTTY is supported by the Singapore National Research Foundation (NRF) Fellowship (Class of 2017). BB acknowledges support from the SickKids Foundation (NI17-039), the National Sciences and Engineering Research Council of Canada (NSERC; Discovery-1304413), CIHR (FDN-154298), Azrieli Center for Autism Research (ACAR), an MNI-Cambridge collaboration grant, and the Canada Research Chairs program. CP was funded through a postdoctoral fellowship of the Fonds de la Recherche due Quebec – Santé (FRQ-S).

## Supplementary Results

### Replication of structural covariance gradients in eNKI dataset

To evaluate whether the observed organizational axes of structural covariance could also be observed in different datasets with a wider age-range, we evaluated the structural covariance gradients in the eNKI dataset (792 individuals, ages 8-85yrs). Here we observed, similar to the main observations in the HCP dataset, a principal anterior posterior gradient explaining 15% of variance and a secondary gradient traversing from inferior to superior regions explaining 11% of variance. Though overall patterns were highly comparable (G1: r_spin_=0.81, p<0.0001, G2: r_spin_=0.88, p<0.0001) between HCP and eNKI covariance gradients (Supplementary Figure 1).

### Association between ageing and structural covariance organization axes

As the eNKI dataset had a broad age distribution we evaluated whether the effect of age on covariance was also organized along posterior-anterior and inferior-superior axis. For this we computed the t-maps of age-related modulation of covariance, and performed gradient analysis on the t-maps. Again, we observed a principal gradient (14% of variance) traversing from posterior to anterior regions, and a secondary gradient (12% of variance) traversing from inferior to superior regions. These gradients showed high correlation with the overall principal and secondary gradients in this dataset (G1: r_spin_=0.76, p<0.0001, G2: r_spin_=0.63, p<0.0001) (Supplementary Figure 3).

### The third – eight gradient of thickness covariance and genetic correlation of thickness

Additionally, we studied the third-eight gradient of thickness covariance and genetic correlation of thickness, explaining 5-10% of variance (Supplementary Figure 1 and 2). The third gradient traversed from sensory-motor and mid temporal areas to both frontal and occipital cortices, and a comparable gradient was observed in genetic correlation of thickness. The fourth gradient had a bilateral axis in superior dorsolateral frontal cortex on the one hand and frontal polar, parietal and temporal polar regions on the other hand. The fifth gradient showed strong lateralization between left temporal parietal regions and right lingual gyrus and corresponded to the sixth gradient of genetic correlation of thickness. The sixth gradient was centered in the right supramarginal gyrus extending to sensory-motor areas on the one hand, and less so in the left sensory cortex, and on the other hand precuneus and para-limbic areas, a similar gradient was not observed in genetic correlation of thickness. The seventh gradient related to sensory-motor, fusiform gyrus and posterior-mid cingulate on the one hand, and temporal regions and precuneus on the other and was most pronounced in the right hemisphere, this gradient was similar to the fifth gradient in coheritability of thickness. The eighth gradient showed a dissociation between temporal parietal regions and posterior-mid cingulate on the one hand, and occipital and sensory regions on the other.

### Structural gradients are above and beyond geodesic distance

Previous work has shown a strong relationship between structural thickness covariance, genetic correlation of cortical thickness, and geodesic distance ^15^. Thus, we explored the relationship between organization of structural covariance and geodesic distance. Geodesic distance was defined as the average distance between each of the 400 parcels ipsilaterally (Supplementary Figure 10). In line with previous reports, we observed a strong relation between structural covariance and geodesic distance (left hemisphere: r=-0.52, p<0.00001, right hemisphere: r=0.51, p<0.00001). Moreover, we observed that genetic correlation varied as a function of the organization of distance, with regions at comparable levels of the geodesic distance gradients showing high genetic correlation among each other. Importantly, when controlling for geodesic distance we again observed an inferior-superior gradient and a posterior-anterior gradient, suggesting the organizational patterns in covariance exist above and beyond geodesic distance. Notably, comparing the topological organization based on geodesic distance and structural covariance, we observed that especially regions in the temporal-parietal areas showed stronger covariance than expected based on distance along, whereas regions in sensory-motor areas showed less covariance than expected based on distance (Supplementary Figure 11).

### Relationship between large-scale organization of genetic correlation of regional thickness and microstructure profiles

In a last step we evaluated the association between the two main axis of regional covariance topology and cortical microstructure (T1w/T2w), microstructural covariance gradients ^6^, and large-scale organization of functional connectivity ^7^, in order to qualify and quantify the relation of the observed covariance gradients in thickness to previously reported microstructural and functional cortical organization^6, 7^. We probed cortical microstructure at 12 equidistant surfaces sampled between the outer and inner cortical layer ^6^ in the same participants (HCP S900 sample). We observed a strong negative relationship between G1_scov_ and cortical T1w/T2w at all layer depts (−0.34 < r >−0.44) (Supplementary Figure 12A; Supplementary Table 4). G2_scov,_ however, only showed a significant positive association with the two most outer strata (layer 1: r=0.60, layer 2: r=0.40), but not with layers closer to the GM/WM surface (Supplementary Figure 12A; Supplementary Table 5). Following we probed the association between organizational gradients of within-individual microstructural profile covariance and topological organization of structural covariance of cortical thickness. To do so, we computed the mean microstructural profile covariance (MPC) maps across individuals and preformed gradient decomposition. We observed, as previously reported ^6^, a primary gradient of cortical microstructural profile covariance traversing a sensory-fugal pattern (22% of variance), and secondary gradient (17% of variance) traversing a pattern from sensory-motor to frontal cortices. We found that the first MPC gradient showed a close correlation with the inferior-superior gradient of genetic covariance of thickness (r=0.62, p<0.00001), but not with the posterior-anterior gradient of genetic covariance of thickness (r=-0.02). Conversely, the secondary gradient of MPC was associated with the posterior-anterior gradient of genetic covariance of thickness (r=0.30, p<0.00001), but not with the inferior-superior gradient of genetic covariance (r=-0.09, p>0.1).

### Relationship between large-scale organization of genetic correlation of regional thickness and functional connectivity topology

Next, we evaluated the association between the posterior-anterior and inferior-superior covariance gradients and the previously reported large-scale organizational gradient of functional connectivity (constructed based on functional connectivity maps in a subset of the HCP S900 sample)^7^ (Supplementary Figure 12). We observed that the functional gradient showed a positive correlation with the rostral-caudal gradient (r=0.37 [0.23 0.49], p<0.00001) but not with the ventral-dorsal gradient alone did not relate to the large-scale functional gradient (r=0.08 [-0.04 0.23], p<0.1). At the same time, the combination of the both gradients showed a strong association with large-scale functional organization (r=0.45 [0.33 0.58], p<0.00001), above and beyond the association with rostro-caudal patterns alone (r_diff_ −0.08 [−0.18 −0.01]). Indeed, combining the rostro-caudal and ventral dorsal gradient partially revealed an organization patterns from unimodal (visual and sensory-motor cortex) to heteromodal association areas (frontal and temporal cortex). Genetic correlation was observed to vary as a function of the combination of gradients and was strongest in regions at similar levels of the combined gradient. Last, we evaluated genetic correlation patterns as a function of the functional gradient reported by Margulies^7^. We observed genetic correlation also varied as a function of large-scale organization of functional connectivity, with regions at similar gradient levels (probed in 10 equally sized bins) showing stronger genetic correlation relative to regions at different gradient levels.

### Functional topography along macro scale organizational patterns of thickness

We conducted a meta-analysis using the Neurosynth ^97^ database and estimated the center of gravity across a set of diverse cognitive terms ^5, 13^ along the posterior-anterior and inferior-superior macro scale organization patterns of thickness (Supplementary Figure 13). In the posterior-anterior gradient we observed a divergence between sensory and visual functions posteriorly and ‘working-memory’, ‘reading’, as well as ‘motor’ and ‘action’ processing anteriorly. Various terms such as ‘emotion’ and ‘reward’ related to both posterior and anterior regions. The inferior-superior gradient on the other hand related to ‘motor’, ‘working memory’ and ‘action’ in superior regions, but ‘emotion’, ‘reward’, ‘affective’, ‘pain’ in inferior regions.

## SUPPLEMENTARY TABLES

**Supplementary Table 1.** Average structural covariance (Spearman’s rho) in each **cytoarchitectural and functional network.** Corresponding table to Figure 1Bi, indicating the average covariance of regions within the respective functional networks.

| Functional networks | Visual | Sensory-Motor | Dorsal attention | Ventral attention | Limbic | Fronto-parietal network | Default mode network |
| --- | --- | --- | --- | --- | --- | --- | --- |
| Visual | 0,09 | -0,01 | -0,01 | -0,02 | -0,01 | -0,04 | -0,03 |
| Sensory Motor | -0,01 | 0,04 | 0,01 | 0,00 | -0,03 | -0,01 | -0,02 |
| Dorsal attention | -0,01 | 0,01 | 0,05 | -0,01 | -0,05 | 0,01 | -0,01 |
| Ventral attention | -0,02 | 0,00 | -0,01 | 0,01 | -0,01 | 0,00 | 0,00 |
| Limbic | -0,01 | -0,03 | -0,05 | -0,01 | 0,09 | -0,01 | 0,01 |
| Fronto-parietal control network | -0,04 | -0,01 | 0,01 | 0,00 | -0,01 | 0,03 | 0,01 |
| Default mode network | -0,03 | -0,02 | -0,01 | 0,00 | 0,01 | 0,01 | 0,02 |

**Supplementary Table 2.** Average genetic correlation between each functional network. Table based on Figure 2Ai.

| Functional networks | Visual | Sensory-Motor | Dorsal attention | Ventral attention | Limbic | Fronto-parietal network | Default mode network |
| --- | --- | --- | --- | --- | --- | --- | --- |
| 1 | 0,25 | 0,00 | 0,00 | -0,06 | -0,01 | -0,11 | -0,08 |
| 2 | 0,00 | 0,10 | 0,04 | 0,00 | -0,07 | -0,04 | -0,05 |
| 3 | 0,00 | 0,04 | 0,20 | 0,02 | -0,15 | 0,04 | -0,02 |
| 4 | -0,06 | -0,01 | 0,01 | 0,04 | -0,03 | 0,01 | 0,01 |
| 5 | -0,01 | -0,07 | -0,14 | -0,03 | 0,20 | -0,03 | 0,02 |
| 6 | -0,11 | -0,04 | 0,04 | 0,01 | -0,02 | 0,10 | 0,05 |
| 7 | -0,08 | -0,05 | -0,02 | 0,01 | 0,02 | 0,06 | 0,05 |

**Supplementary Table 3.**
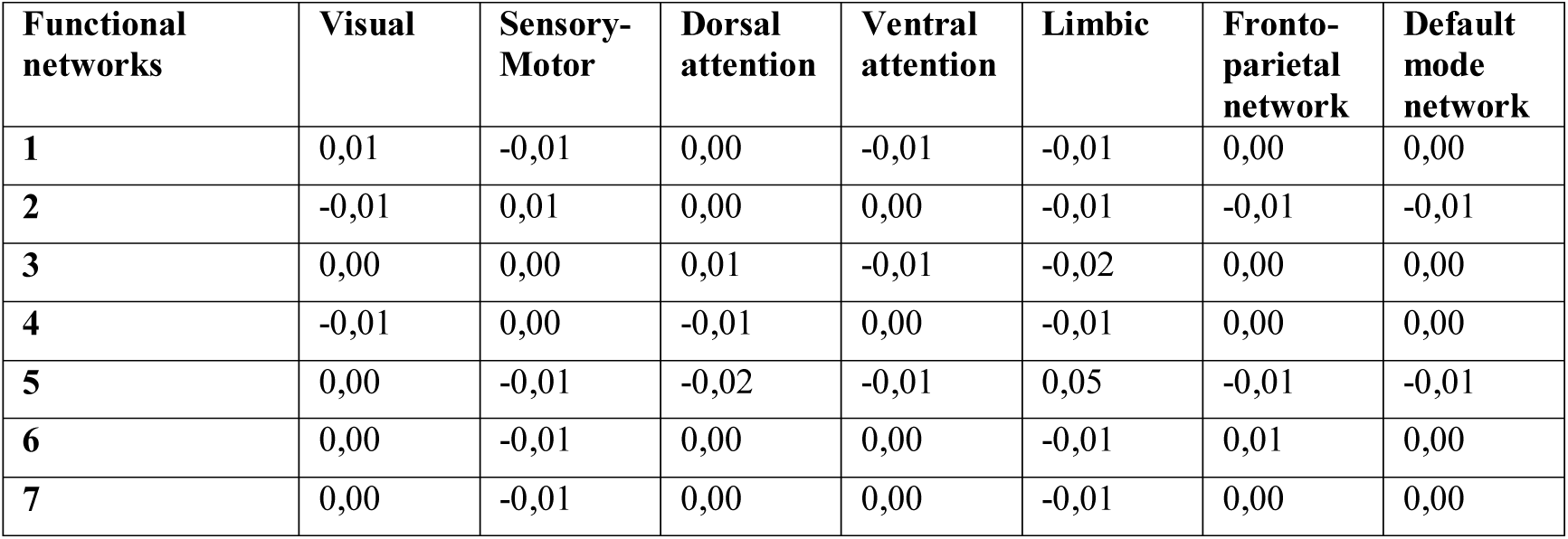
Average environmental correlation between each functional **network.** Table based on Figure 2Bi.

**Supplementary Table 4.** Correlation between layer-dependent T1q and G1_GC_. Correlation between layer-based T1w/T2w and the primary gradient of thickness covariance.

| T1w/T2w | Correlation with G1 | T1w/T2w | Correlation with G1 |
| --- | --- | --- | --- |
| Layer 1 | -0.34, $p < 0.000001$ | Layer 7 | -0.42, $p < 0.000001$ |
| Layer 2 | -0.40, $p < 0.000001$ | Layer 8 | -0.43, $p < 0.000001$ |

|  |  |  |  |
| --- | --- | --- | --- |
| Layer 3 | -0.40, $p < 0.000001$ | Layer 9 | -0.43, $p < 0.000001$ |
| Layer 4 | -0.38, $p < 0.000001$ | Layer 10 | -0.44, $p < 0.000001$ |
| Layer 5 | -0.39, $p < 0.000001$ | Layer 11 | -0.43, $p < 0.000001$ |
| Layer 6 | -0.40, $p < 0.000001$ | Layer 12 | -0.43, $p < 0.000001$ |

**Supplementary Table 5.** Correlation between layer-dependent T1q and G2_GC_. Correlation between layer-based T1w/T2w and the secondary gradient of thickness covariance.

| T1w/T2w | Correlation with G2 | T1w/T2w | Correlation with G2 |
| --- | --- | --- | --- |
| Layer 1 | 0.64, $p < 0.000001$ | Layer 7 | -0.05, $p > ns$ |
| Layer 2 | 0.40, $p < 0.000001$ | Layer 8 | -0.04, $p > ns$ |
| Layer 3 | 0.12, $p < 0.02$ | Layer 9 | -0.03, $p > ns$ |
| Layer 4 | -0.01, $p > ns$ | Layer 10 | -0.03, $p > ns$ |
| Layer 5 | -0.05, $p > ns$ | Layer 11 | -0.02, $p > ns$ |
| Layer 6 | -0.05, $p > ns$ | Layer 12 | 0.00, $p > ns$ |

## SUPPLEMENTARY FIGURES

**Supplementary Fig 1.**
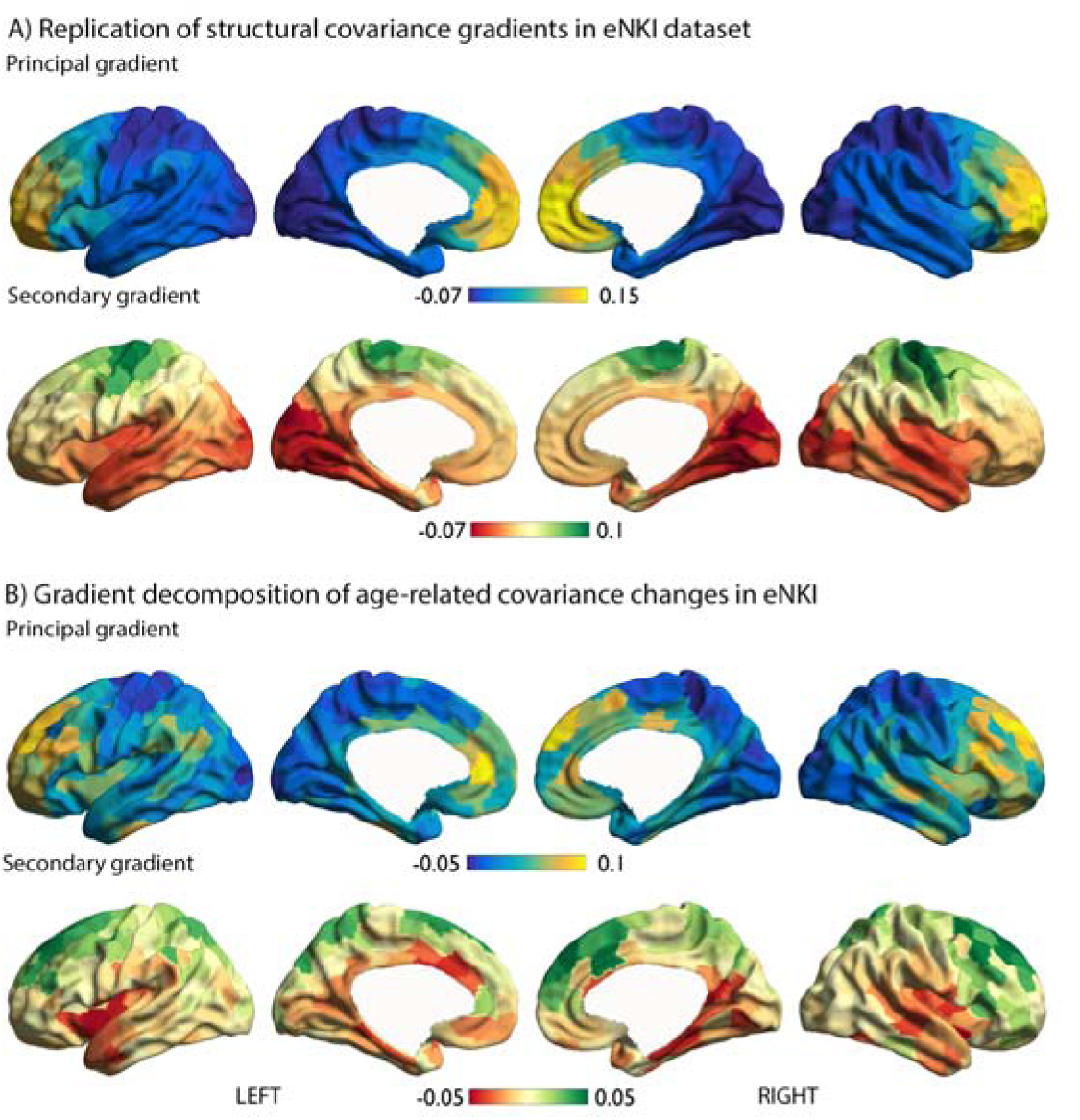
Robustness of structural covariance gradients using replication sample (eNKI) and associations with age-related change in covariance. A). Replication of the first two gradients in the eNKI dataset, using the Schaefer 400 parcellation. B). Gradient decomposition of t-maps of age-related modulation of structural covariance.

**Supplementary Fig 2.**
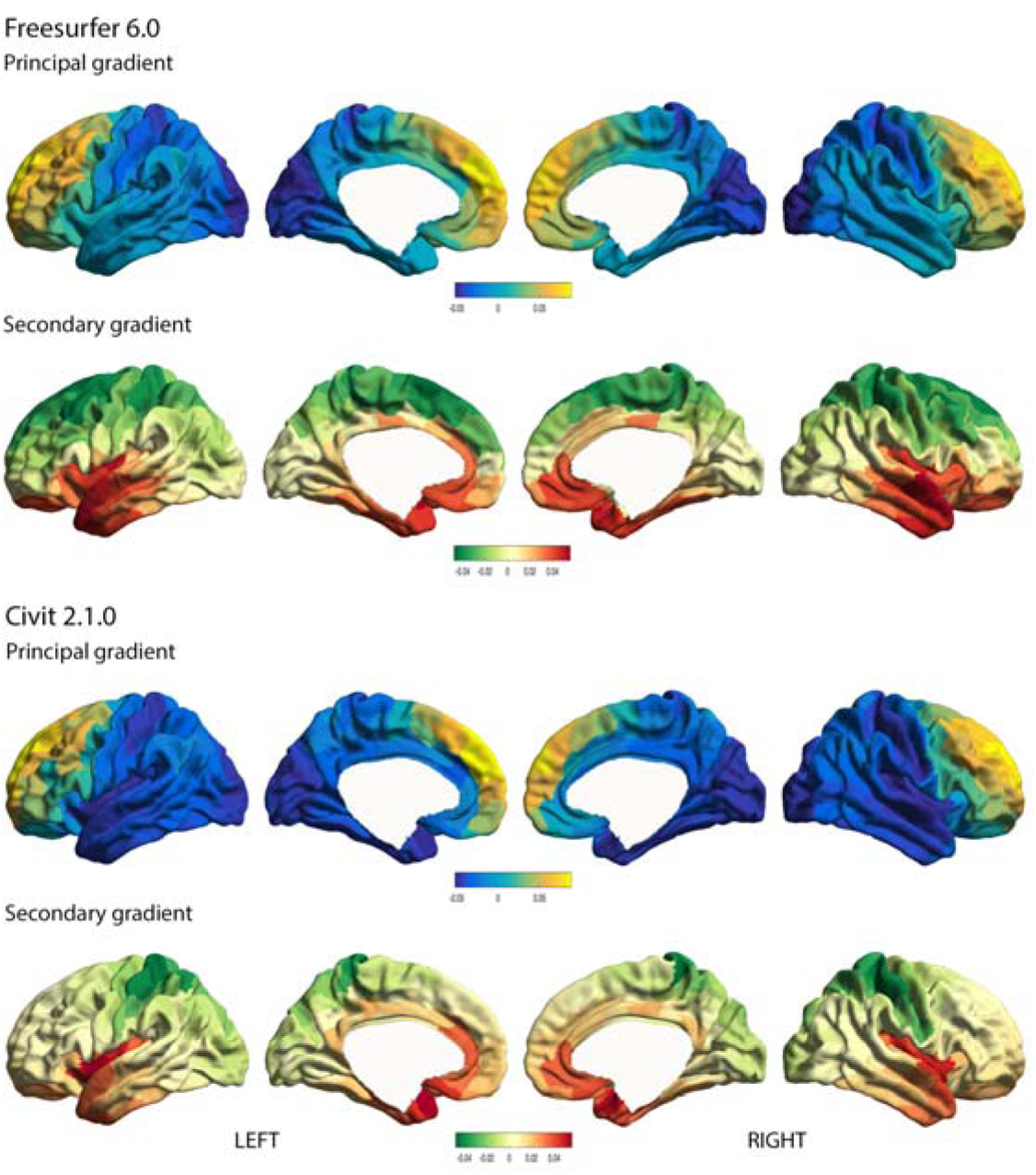
Robustness of structural covariance gradients as a function of cortical thickness estimation method. A). Cortical thickness estimation in HCP sample based on Freesurfer 6.0 standard pipeline. B). Cortical thickness estimation in HCP sample based on CIVIT 2.1.0. standard pipeline.

**Supplementary Figure 3.**
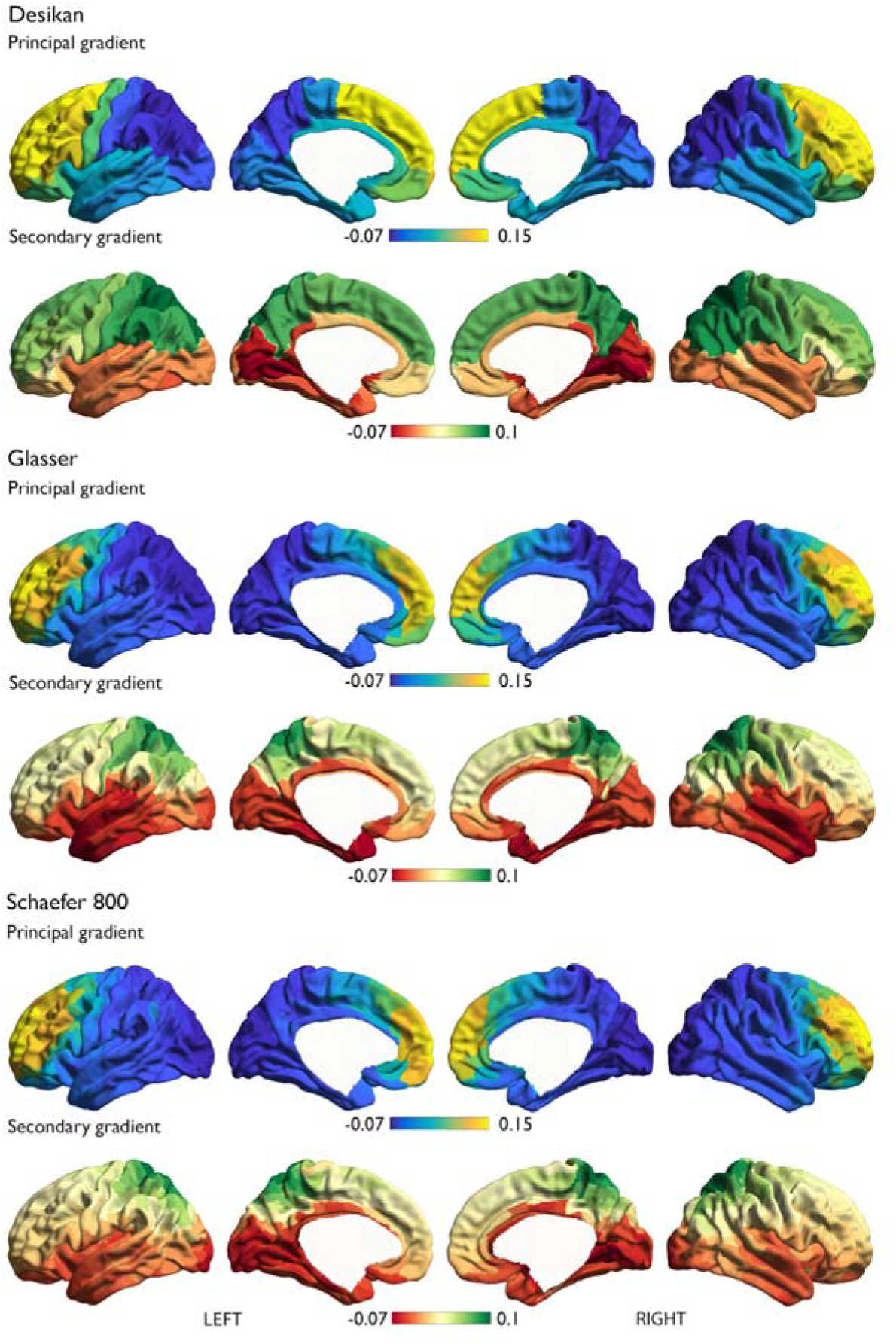
Robustness of structural covariance gradients as a function of parcellation method. A). Cortical thickness parcellated using the Desikan-Killiany atlas; B). Cortical thickness parcellated using the Glasser atlas; C). Cortical thickness parcellated using the Schaefer 800 atlas

**Supplementary Figure 4.**
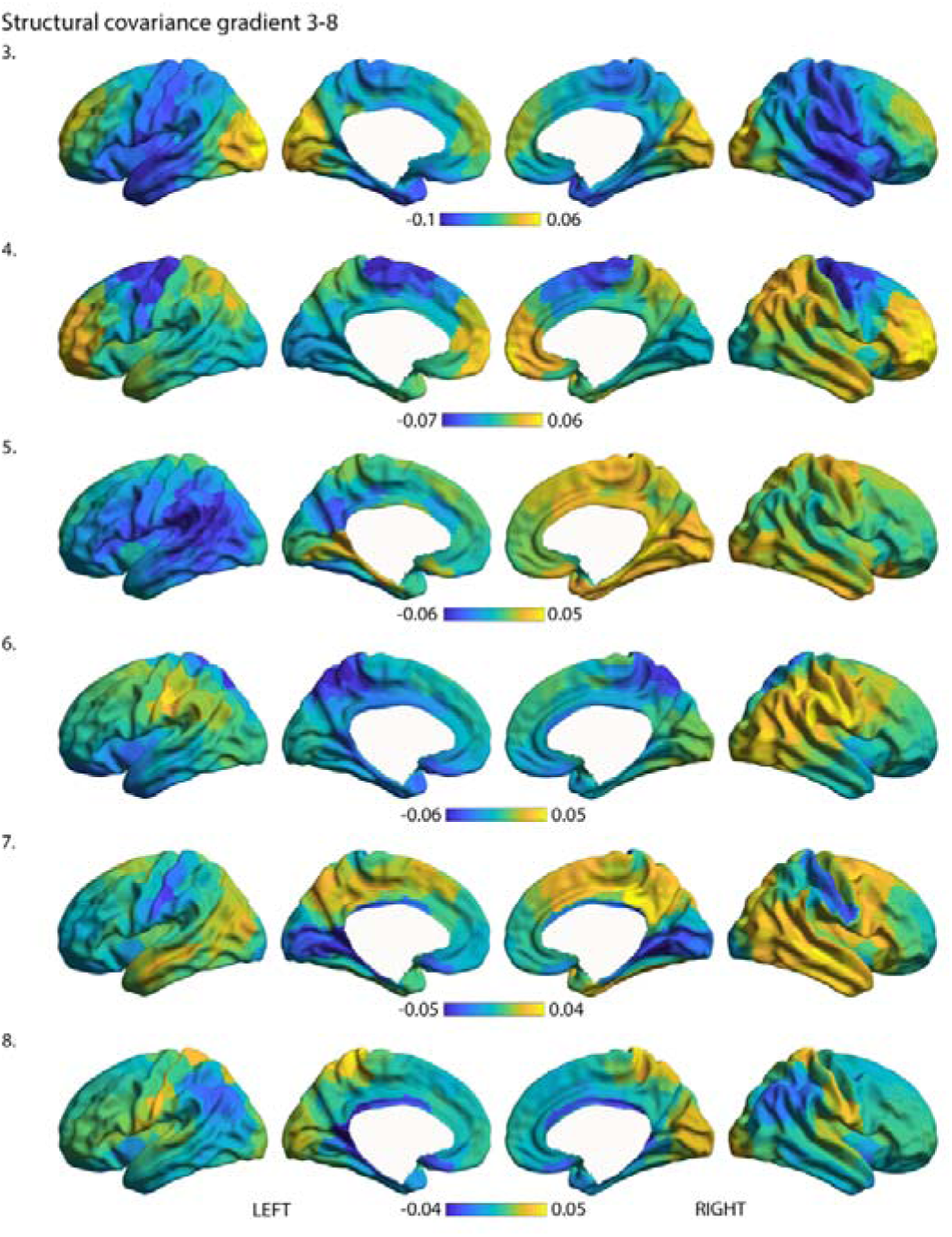
Structural covariance gradients 3-8. The third-eight gradient of structural covariance of thickness.

**Supplementary Figure 5.**
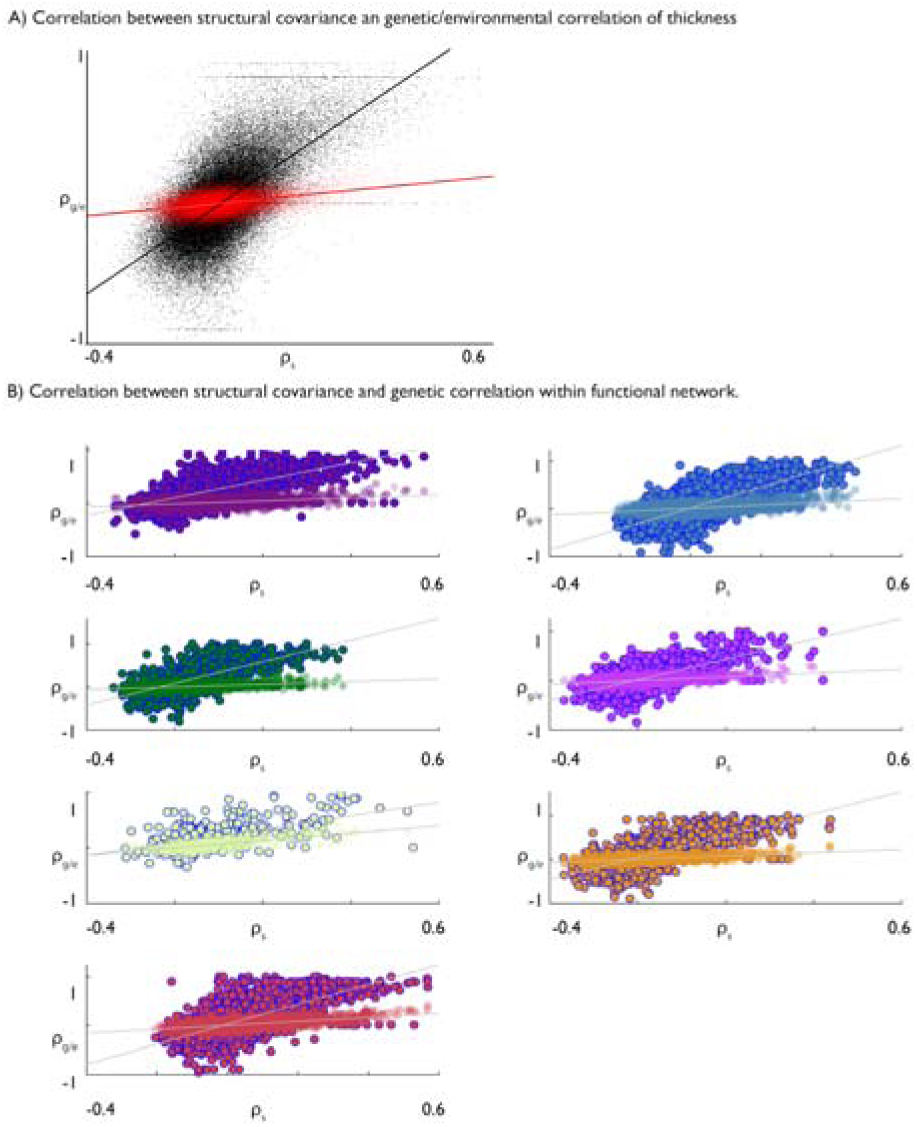
Correlation between structural covariance of thickness and genetic and environmental components. A). Whole brain correlation between covariance and genetic correlation (black) and environmental correlation (red). B). Correlation between covariance and genetic correlation (blue outline) and environmental correlation (no outline) within each functional community^25^.

**Supplementary Figure 6.**
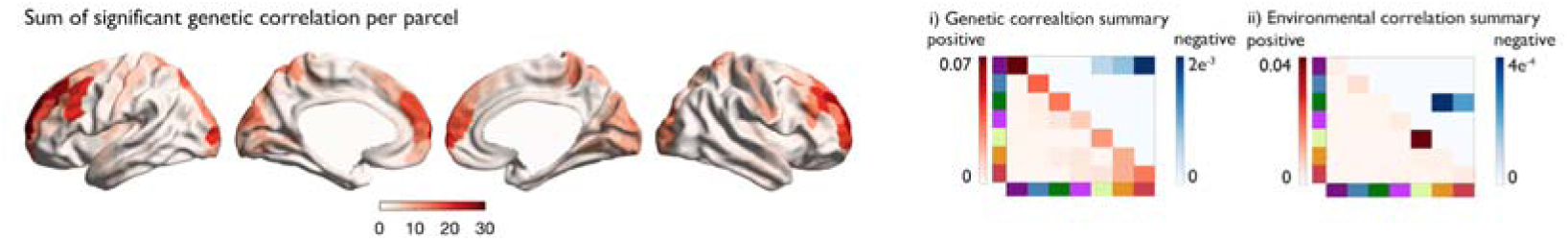
Spatial distribution of significant genetic correlations. Sum of significant genetic correlation per parcel (FDRq<0.05); i). genetic correlation summary per functional community (averaged by the total number of parcels in each functional network) (positive: red; negative: blue); ii). genetic correlation summary per functional community (averaged by the total number of parcels in each functional network) (positive: red; negative: blue)

**Supplementary Figure 7.**
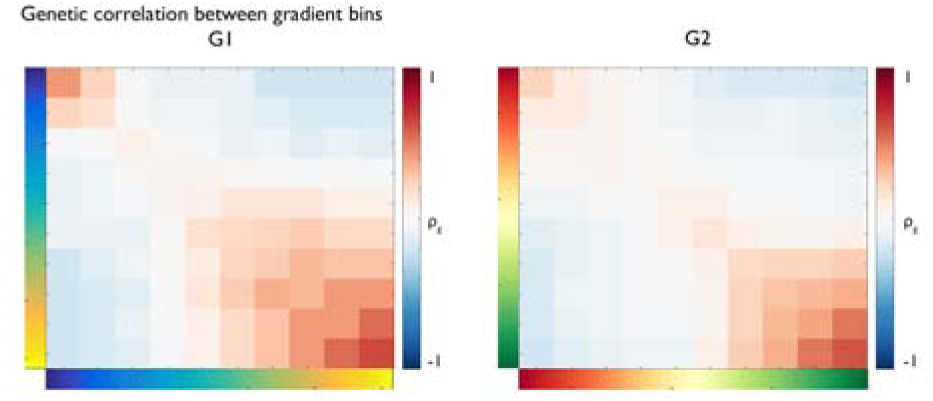
Genetic correlation between gradient bins. The average genetic correlation between binned (10 equally sized bins) principal and secondary gradients of genetic correlation (Figure 2).

**Supplementary Figure 8.**
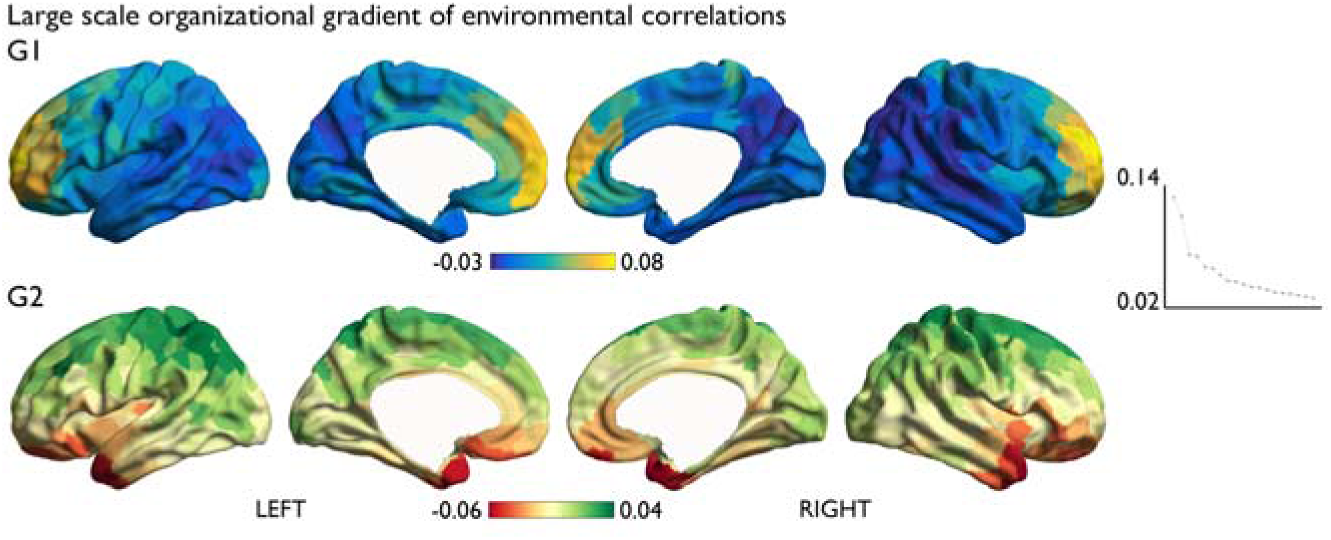
Large-scale organizational gradients of environmental correlations of thickness. Performing the same analysis as in Figure 2C on the environmental correlation of thickness.

**Supplementary Figure 9.**
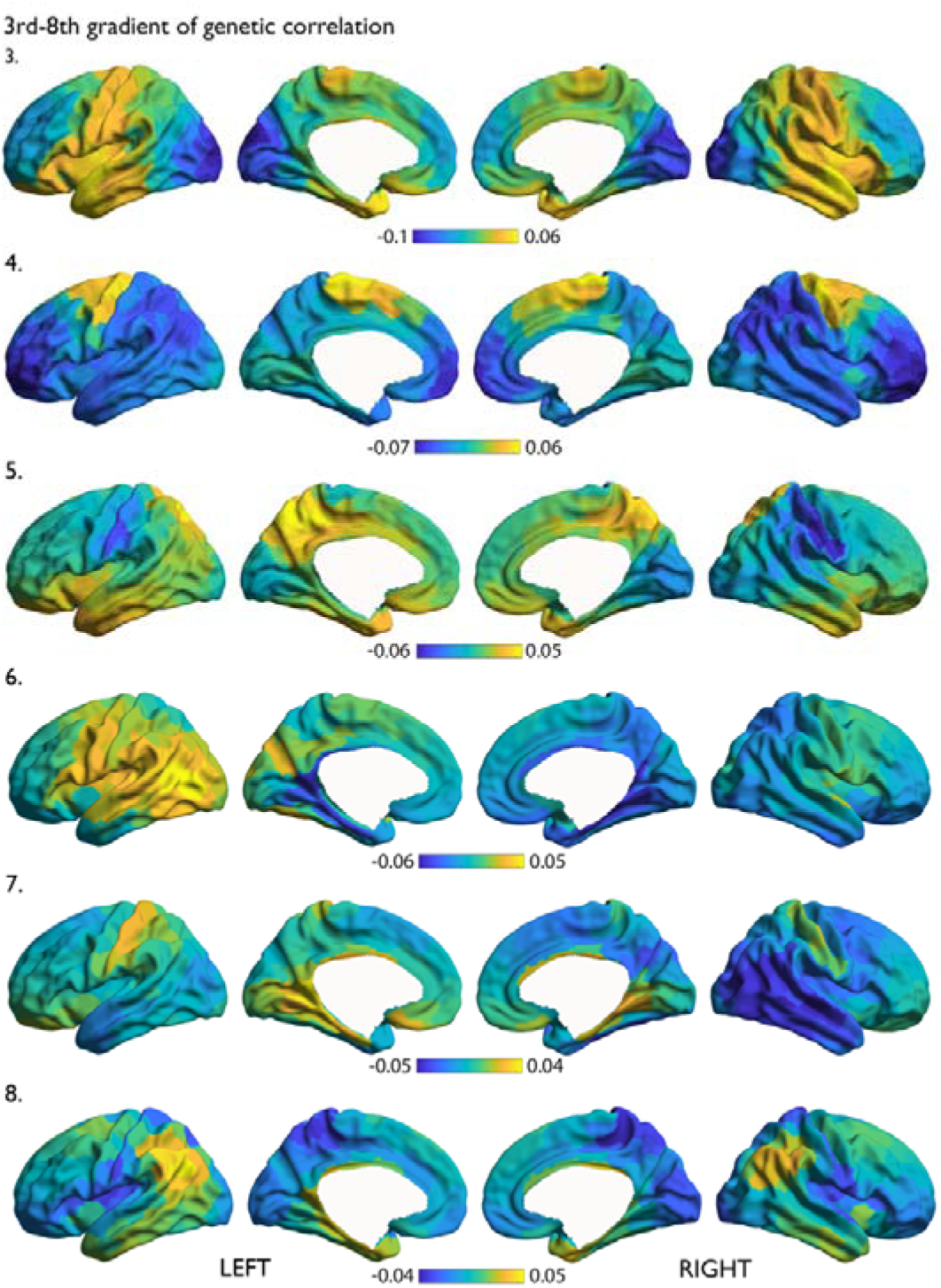
Genetic correlation gradients 3-8. The third-eight gradient of genetic correlation of thickness.

**Supplementary Fig 10.**
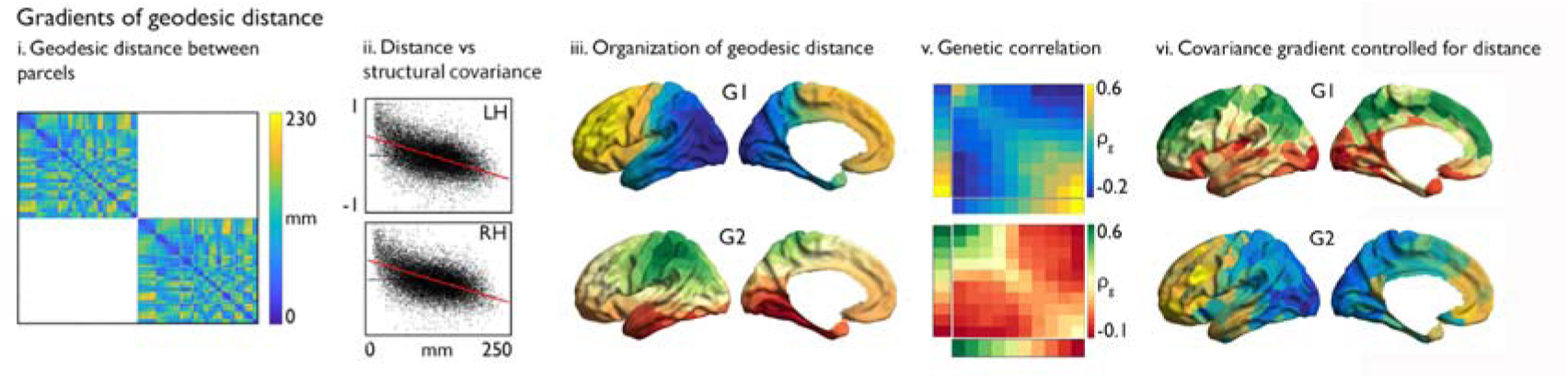
Association between large-scale organization of structural covariance and geodesic distance. i). Geodesic distance matrix of ipsilateral 400 Schaefer parcels; ii). Correlation between geodesic distance and structural covariance between parcels; iii). Principal and secondary gradient of geodesic distance; iv. Genetic correlation as a function of the binned geodesic distance gradients; v. Covariance gradients while controlling for geodesic distance.

**Supplementary Fig 11.**
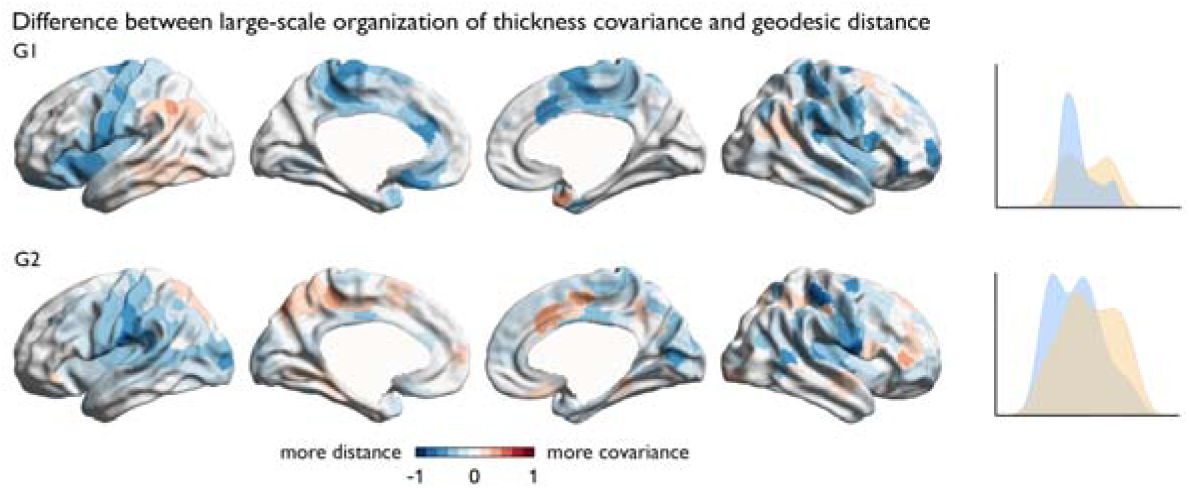
Parcel-wise difference between large-scale organization of structural covariance and geodesic distance. Parcel-wise difference between the structural covariance gradients (G_SCOV_) and the distance-based gradients (G_DIST_). Blue indicates higher gradient ranking in G_DIST_, red indicates higher gradient ranking in G_SCOV_.

**Supplementary Fig 12.**
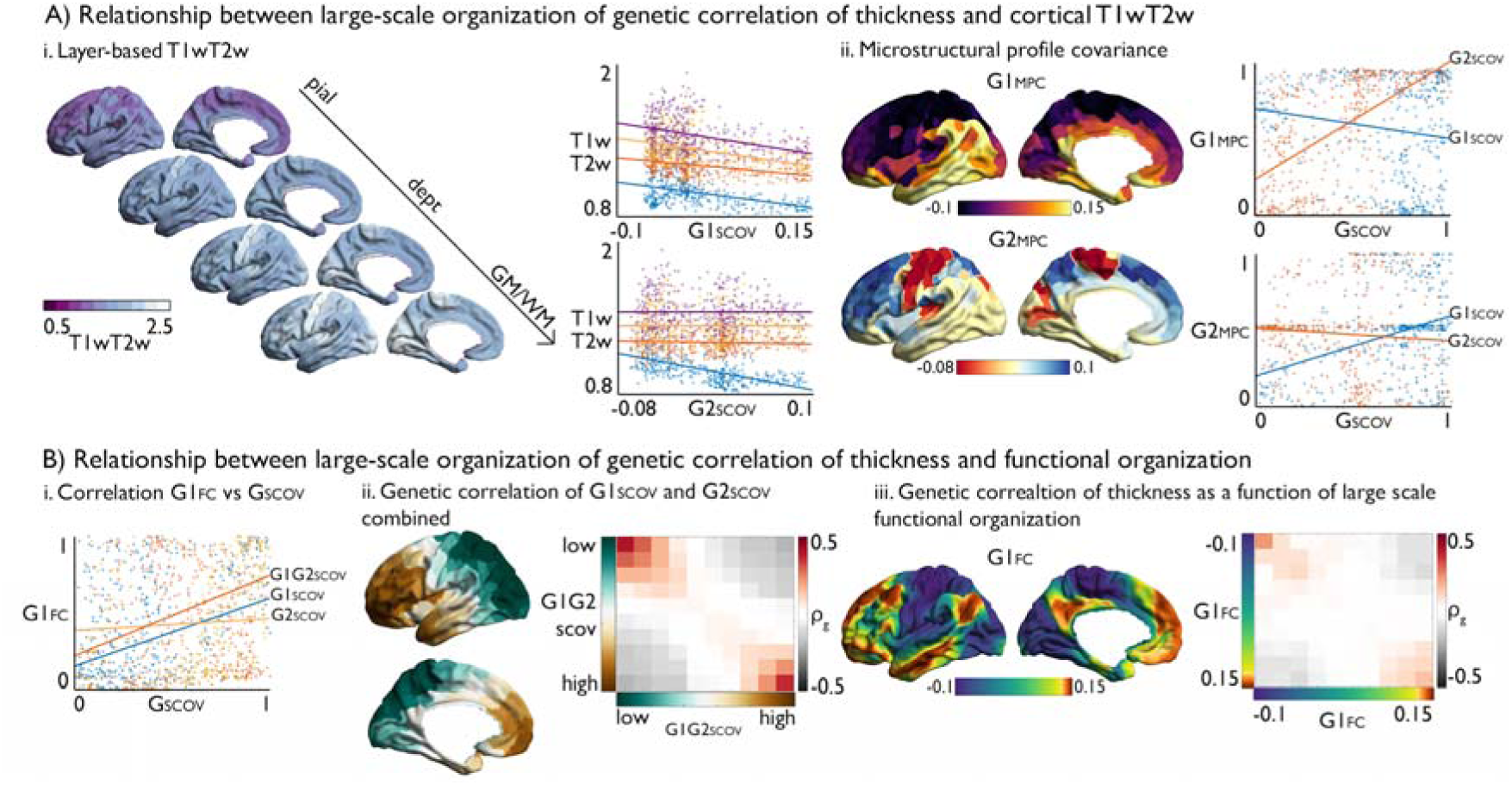
Link between organization of macro scale organization of thickness, microstructure, and function. **A**). Relationship between large-scale organization of genetic correlation of thickness and cortical T1w/T2w; i. T1w/T2w values of equidistant layers between the pial and GM/WM surface and the correlation with the principal and secondary gradient (G1_SCOV_ and G2_SCOV_) of macro scale organization of thickness. For visualization purposes only the first (blue), fourth(orange), seventh (yellow), tenth (purple) of 12 probed layers are reported; ii. Principal and secondary gradient of microstructure profile covariance (MPC) and the relationship between MPC gradients and G1_SCOV_ and G2_SCOV._ **B**). Relationship between large-scale organization of thickness covariance and functional organization; i. the correlation between G1 _SCOV,_ G2 _SCOV_, G1G2_SCOV_ and G1_FC_; ii. Combined G1 _SCOV_ and G2 _SCOV_ gradient, the genetic correlation between binned G1G2 _SCOV_ gradient, and the correlation between G1 _SCOV,_ G2 _SCOV_, G1G2 _SCOV_ and G1_FC;_ ii. Principal gradient of large-scale functional organization and genetic correlation of thickness between G1_FC_ gradient bins.

**Supplementary Fig 13.**
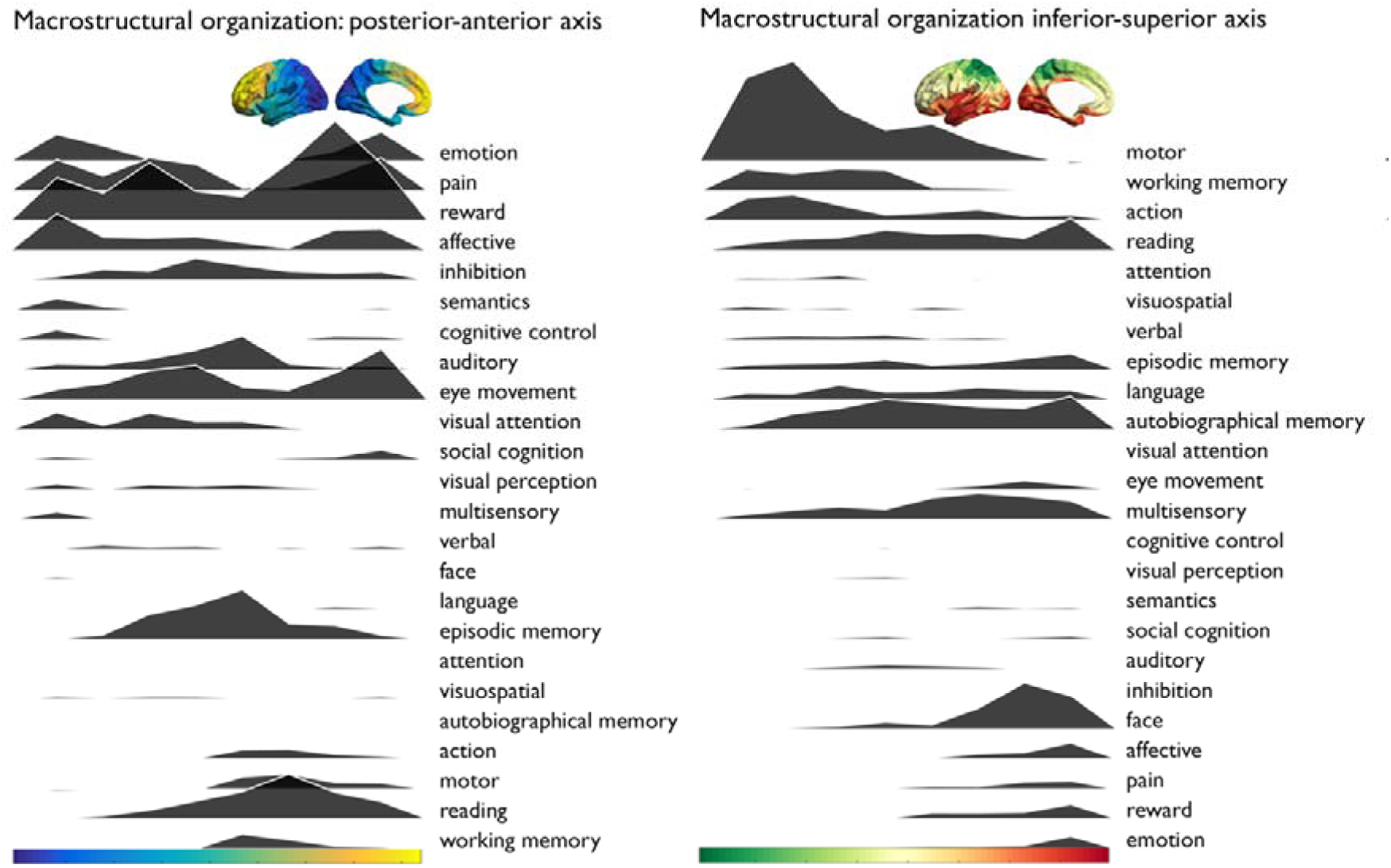
Meta-analysis maps for diverse cognitive terms were obtained from Neurosynth similar to Margulies et al.^13^. We calculated parcel-wise z-statistics, capturing node-term associations, and calculated the center of gravity of each term along the poster-anterior and inferior-superior gradients. The plots depict the average z-score within binned (20-bins) gradient layer of meta-analysis maps.

